# Cryo-EM structures and functional properties of CALHM channels of the human placenta

**DOI:** 10.1101/2020.02.11.944231

**Authors:** Katarzyna Drożdżyk, Marta Sawicka, Maria-Isabel Bahamonde Santos, Jonas Zaugg, Dawid Deneka, Christiane Albrecht, Raimund Dutzler

**Author notes:** These authors contributed equally to this work (Katarzyna Drożdżyk, Marta Sawicka). For correspondence (RD), (MS).

## Abstract

The transport of substances across the placenta is essential for the development of the fetus. Here, we were interested in the role of channels of the calcium homeostasis modulator (CALHM) family in the human placenta. By transcript analysis, we found the paralogs CALHM2, 4, and 6 to be highly expressed in this organ and upregulated during trophoblast differentiation. Based on electrophysiology, we found that activation of these paralogs differs from the voltage- and calcium-gated channel CALHM1. Cryo-EM structures of CALHM4 display decameric and undecameric assemblies with large cylindrical pore, while in CALHM6 a conformational change has converted the pore shape into a conus that narrows at the intracellular side, thus describing distinct functional states of the channel. The pore geometry alters the distribution of lipids, which occupy the cylindrical pore of CALHM4 in a bilayer-like arrangement whereas they have redistributed in the conical pore of CALHM6 with potential functional consequences.

## Introduction

The placenta, a complex organ that develops during pregnancy, serves as a hub for the exchange of nutrients and waste products between the mother and the fetus. The flow of substances across the placenta is controlled by two distinct cell-layers, namely fetal capillary endothelial cells and syncytiotrophoblasts (STB). While endothelial cells lining the fetal vessels allow to a certain extent diffusion of (small) molecules through paracellular pathways (Edwards et al., 1993; Lewis et al., 2013), the STB layer constitutes a tight diffusion barrier. STB are polarized multinucleated cells presenting their apical side towards the maternal blood and their basal side towards the fetal capillaries. Hence, the targeted expression of receptors, channels or transport proteins of the STB determines the directionality of transport across the placenta, which is essential for the adequate development of the fetus (Lager and Powell, 2012). Among the transport proteins in the placenta, the role of the calcium homeostasis modulator (CALHM) family is currently unknown. In humans, the CALHM family encompasses six paralogs, some of which function as non-selective channels that are permeable to large substances such as ATP (Ma et al., 2016). CALHM1, the currently best characterized paralog, forms ion channels that are activated by depolarization and a decrease in the extracellular Ca^2+^-concentration (Ma et al., 2012; Taruno et al., 2013b) with mutations being associated with an increased risk for late-onset Alzheimer’s disease (Dreses-Werringloer et al., 2008). Although functional on its own (Ma et al., 2012), in a physiological context CALHM1 probably forms heteromers with the subunit CALHM3 (Ma et al., 2018b). These heteromeric channels play an important role as secondary receptors for sweet, bitter and umami taste reception in type II taste bud cells by catalyzing the non-vesicular release of ATP (Ma et al., 2018b; Taruno et al., 2013a; Taruno et al., 2013b). A second paralog with an assigned physiological role is CALHM2, which is ubiquitously expressed and was proposed to mediate ATP release in astrocytes (Ma et al., 2018a). The role of other paralogs has thus far remained elusive. Insight into the structural properties of the family was recently provided from studies on CALHM1 and 2 by cryo-electron microscopy (Choi et al., 2019; Demura et al., 2020; Syrjanen et al., 2020). CALHM2 was found to assemble into undecameric channels, which in certain cases were found to dimerize via contacts on the extracellular side in an assembly that resembles gap-junctions. Based on this architecture, a potential role of CALHM2 as intercellular channels was proposed. Although predicted to be related to connexins and volume-regulated anion channels (VRACs) of the LRRC8 family (Siebert et al., 2013), the four membrane-spanning helices of the CALHM subunit exhibit a distinct arrangement, which refutes a common structural ancestry between these protein families (Choi et al., 2019). In contrast to CALHM2 channels, the structure of CALHM1 shows a smaller octameric organization (Demura et al., 2020; Syrjanen et al., 2020), thus suggesting that the functional properties of activation and conduction might not be conserved within the family.

In the present study we were investigating the structural and functional properties of CALHM channels in the context of the placenta. We identified the three paralogs, CALHM2, 4 and 6, to be highly expressed in this organ. A systematic comparative functional characterization by electrophysiology did not reveal pronounced activity of either of these paralogs under conditions where CALHM1 channels are open, thus suggesting that the former might be regulated by different and currently unknown mechanisms. The structural characterization of the three paralogs reveals insight into their organization and into potential gating transitions which, although related to previously described properties of CALHM2, show distinct features with respect to subunit organization, conformational changes and the distribution of lipids residing inside the wide pore of CALHM channels.

## Results

### Expression of CALHM channels in the human placenta

To investigate the role of CALHM channels in the placenta, we have characterized the expression of family members in healthy human placental tissues obtained from term pregnancies. Quantification of transcripts by reverse transcription (RT) PCR revealed high levels of CALHM2, 4, and 6, but comparably low expression of other paralogs (Figure 1A). We also analyzed the expression patterns of CALHM paralogs in primary trophoblast cells isolated from healthy term placentae and examined changes during the differentiation process of trophoblast precursor cells into mature STB. We found marked upregulation of CALHM2, 4 and 6 during differentiation, which was most pronounced for CALHM4 (Figure 1B) and which might indicate a differentiation-dependent role of these proteins in the human placenta. In all cases, the concentrations of the respective CALHM mRNAs are high in comparison to other transport proteins (Figure 1-figure supplement 1) suggesting their involvement in important placenta-related membrane transport processes.

**Figure 1.**
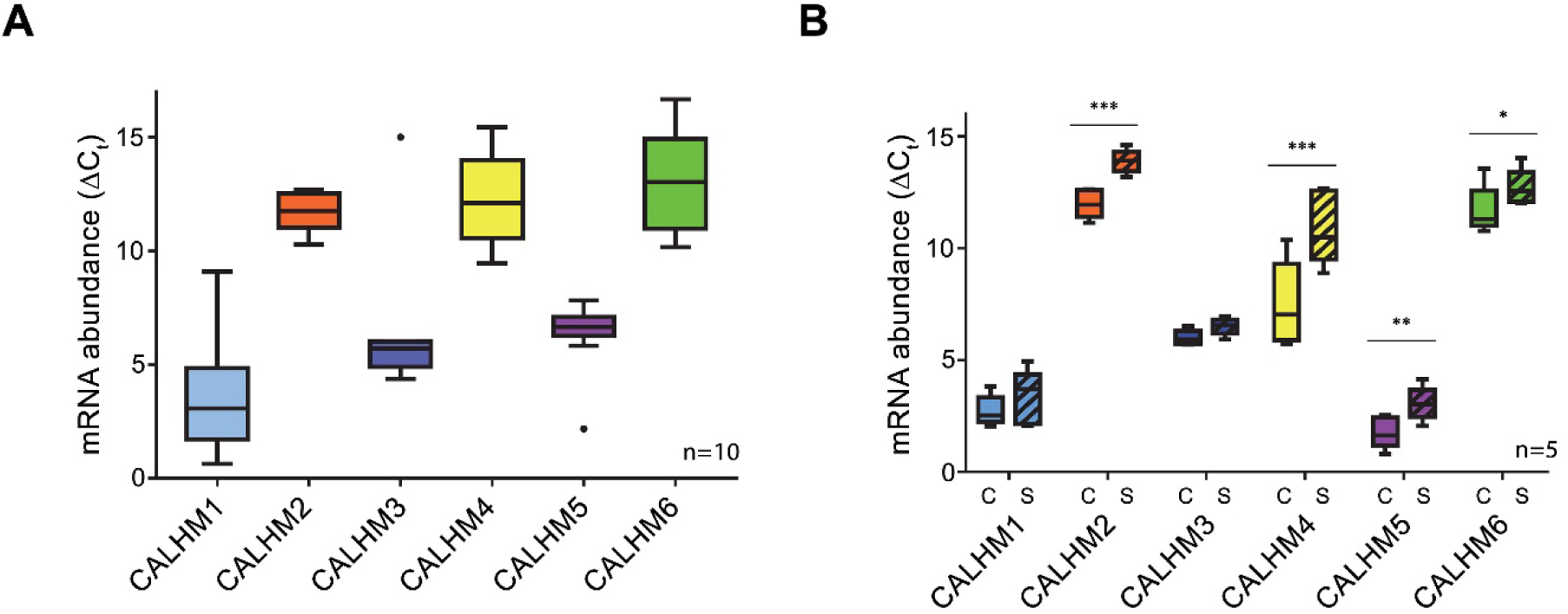
Expression analysis of CALHM genes in the human placenta. mRNA abundances of different CALMH paralogs in (**A**), human placental tissues (n=10) and (**B**), human trophoblasts isolated from healthy term placentae (n=5) was assessed by quantitative RT PCR and normalized to the reference gene YWHAZ. The relative amounts of the different CALHM genes are shown as ΔC_t_ values (ΔC_t_ = C_t_ value of YWHAZ – C_t_ value of CALHM gene). Comparative transcript data are presented as mRNA abundance (ΔCt = Ct_reference gene_ – Ct_traget gene_) between undifferentiated cytotrophoblasts (C) and differentiated syncytiotrophoblast (S) cells. Data analysis and statistical evaluations were performed using paired 2-way ANOVA with Sidak’s multiple comparisons test; * p< 0.05; ** p< 0.01; *** p<0.001.

### Functional characterization of CALHM paralogs

Due to the abundance of CALHM2, 4, and 6 in the placenta, we have focused our subsequent investigations on these three paralogs. Apart from CALHM2, which was described to form ATP-conducting channels in astrocytes (Ma et al., 2018a), they have so far not been characterized. For a functional comparison of channels composed of the three placental subunits with the well-studied protein CALHM1, we have expressed the homomeric proteins in *X. laevis* oocytes and recorded currents by two-electrode voltage-clamp electrophysiology. This method has previously allowed a detailed characterization of CALHM1, which is activated by depolarization and the removal of Ca^2+^ from the extracellular side (Ma et al., 2012). To detect proteins in the plasma membrane, we have used a surface-biotinylation approach and found all placental paralogs to be expressed and targeted at high levels to the surface of oocytes at the time of the measurement (*i.e.* 42-52 hours after injection of mRNA) (Figure 2A). Assuming that CALHM proteins form channels of large conductance, we thus can directly compare the average magnitude of recorded currents between populations of oocytes expressing the respective constructs and relate their activation properties in response to voltage change and the depletion of extracellular Ca^2+^ to the well-characterized CALHM1. In our studies, we were able to reproduce the functional hallmarks of CALHM1, which are manifested in low currents at negative voltage, slowly activating currents at positive voltage in the presence of extracellular Ca^2+^ and a strong increase of currents upon its removal in the entire voltage range (Figure 2, Figure 2-figure supplement 1A and B). In contrast to CALHM1, the current response of homomeric CALHM2, 4, and 6 channels was generally small and within the range of endogenous currents of *X. laevis* oocytes not expressing any of the proteins (Figure 2A, Figure, 2-figure supplement 1C-F). These currents neither showed pronounced voltage-dependence nor were they altered by Ca^2+^-removal in a statistically significant manner (Figure 2B-D). We thus conclude that the CALHM channels expressed in the placenta are not regulated in a similar manner as CALHM1-subunit containing channels and that their activation instead proceeds by distinct, currently unknown mechanisms.

**Figure 2.**
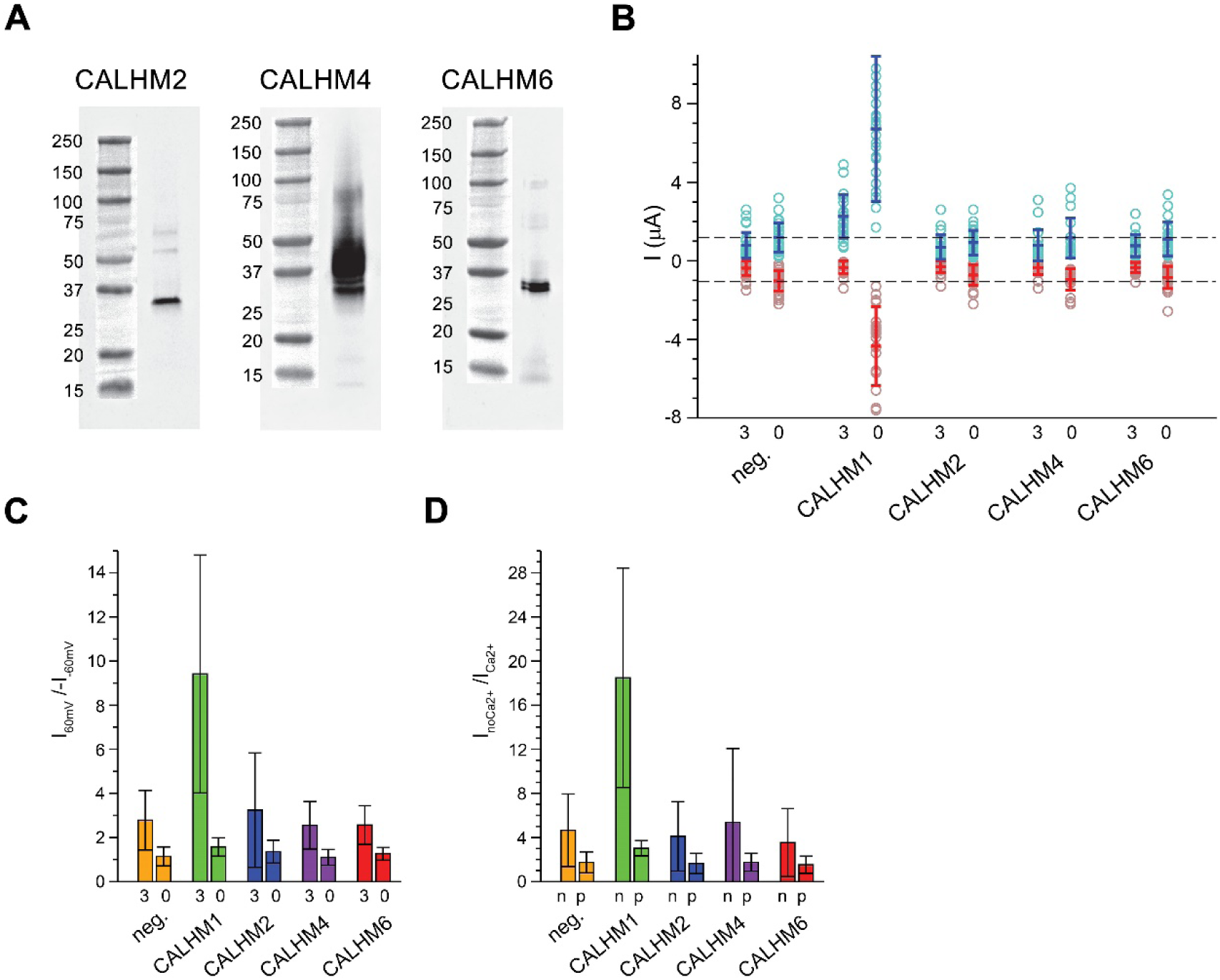
Functional characterization of CALHM channels expressed in *X. laevis* oocytes. (**A**) Western blot of proteins located in the plasma membrane of *X. laevis* oocytes heterologously expressing the indicated CALHM channels. Protein was isolated after surface-biotinylation by binding to avidin-resin. The proteins are detected with specific antibodies recognizing the respective CALHM paralog. Left, CALHM2, center CALHM4, right CALHM6. The blot demonstrates the targeting of all three paralogs to the plasma membrane. (**B**) Electrophysiological characterization of *X. laevis* oocytes heterologously expressing the paralogs CALHM1, CALHM2, CALHM4 and CALHM6 in comparison to control oocytes (neg.) recorded at extracellular solutions either containing 3 mM Ca^2+^ (3) or 0.5 mM EDTA (Ca^2+^-free, 0). Data show currents of individual oocytes (circle) recorded by two-electrode voltage-clamp (TEVC) at 60 (light blue) and −60 mV (light red). Averages are shown as bars in red (−60 mV) and blue (60 mV), respectively. Dashed lines indicate mean current levels of control oocytes (neg.) recorded in Ca^2+^-free extracellular solutions at 60 and −60 mV. (**C**), Rectification of steady-currents of oocytes displayed in (B) expressed as I_60mV_/I_−60mV_ calculated for individual oocytes at 3 mM Ca^2+^ (3) and in Ca^2+^-free solutions (0) and averaged. The large value of CALHM1 reflects the activation of the protein at positive voltage in presence of Ca^2+^. (**D**) Ca^2+^-dependence of activation. Change of steady-state currents of oocytes displayed in (B) after Ca^2+^-removal expressed as I_noCa2+_/I_Ca2+_ calculated from individual oocytes at −60 mV (n) and 60 mV (p) and averaged. The large value of CALHM1 at −60 mV reflects the strong activation of currents at negative voltages upon Ca^2+^-depletion. C, D, The difference between the corresponding values of the CALHM paralogs 2, 4, and 6 and neg. are statistically insignificant (as judged by a Student t-test). B-D, Data show averages of 27 (neg.), 21 (CALHM1), 26 (CALHM2), 19 (CALHM4) and 21 (CALHM6) oocytes respectively. Errors are standard deviations.

### Biochemical characterization and structure determination of CALHM2, 4, and 6

For a biochemical and structural characterization of placental CALHM channels, we have expressed constructs coding for human CALHM2, 4 and 6 in HEK293 cells. In contrast to the poor yields obtained for human CALHM1, all three placental paralogs expressed high levels of protein. The elution properties of fusions to green fluorescent protein (GFP) extracted in the detergent glycol-diosgenin (GDN) and analyzed by fluorescent size-exclusion chromatography (FSEC) (Kawate and Gouaux, 2006) indicate assemblies of high molecular weight (Figure 3-figure supplement 1A). We next proceeded with a scale-up of homomeric CALHM2, 4 and 6 channels in HEK293 cells and purified each protein in the detergent GDN. Consistent with FSEC studies, all purified constructs eluted as large oligomers during gel-filtration chromatography, although at different volumes. The highest elution volume was observed for CALHM6, a similar but slightly lower volume for CALHM2 and the smallest volume for CALHM4, thus hinting towards distinct oligomeric organizations of the three proteins with CALHM4 forming larger complexes than CALHM6 (Figure 3-figure supplement 1B). The peak fractions were concentrated, vitrified on cryo-EM grids and used for data collection by cryo-electron microscopy (Figure 3-figure supplements 2-6). As expected from the size-exclusion profiles, all paralogs form large and heterogenic multimeric assemblies containing between 10 and 12 subunits. Whereas in the CALHM6 sample a vast majority of these oligomeric channels do not interact (Figure 3B, Figure3-figure supplement 4), we have found almost complete dimerization for CALHM4 (Figure 3A, Figure 3-figure supplements 2-3) and a significant fraction of particles to dimerize in case of CALHM2 (Figure 3C, Figure3-figure supplement 6). Unlike to previous reports of CALHM2 structures, where interactions between channels were mediated by extracellular loops (Choi et al., 2019; Syrjanen et al., 2020), the pairing in the CALHM4 sample proceeds via contacts at the intracellular side (Figure 3A). We also observed two distinct protein conformations in our data with CALHM4 and CALHM6 channels forming wide cylindrical and conical pores, respectively (Figure 3D). Based on 2D class averages, we assume both conformations to be present in our CALHM2 sample, although 3D classification of this dataset did not yield high-resolution structures. We believe that strong preferential orientation of CALHM2 particles resulting in a predominance of views along the pore axis combined with sample heterogeneity due to the presence of paired and non-paired assemblies of multimeric subunits in our data limited our data processing workflow to 2D classification (Figure 3-figure supplement 6). In general, we found an oligomeric distribution of CALHM2 channels that corresponds to previously determined structure with a majority of channels being organized as undecamers and a smaller population showing dodecameric assemblies (Figure 3C, Figure 3-figure supplement 6). Due to the higher quality of the CALHM4 and 6 samples, we continued to use structures derived from these proteins for a detailed characterization of both pore conformations.

**Figure 3.**
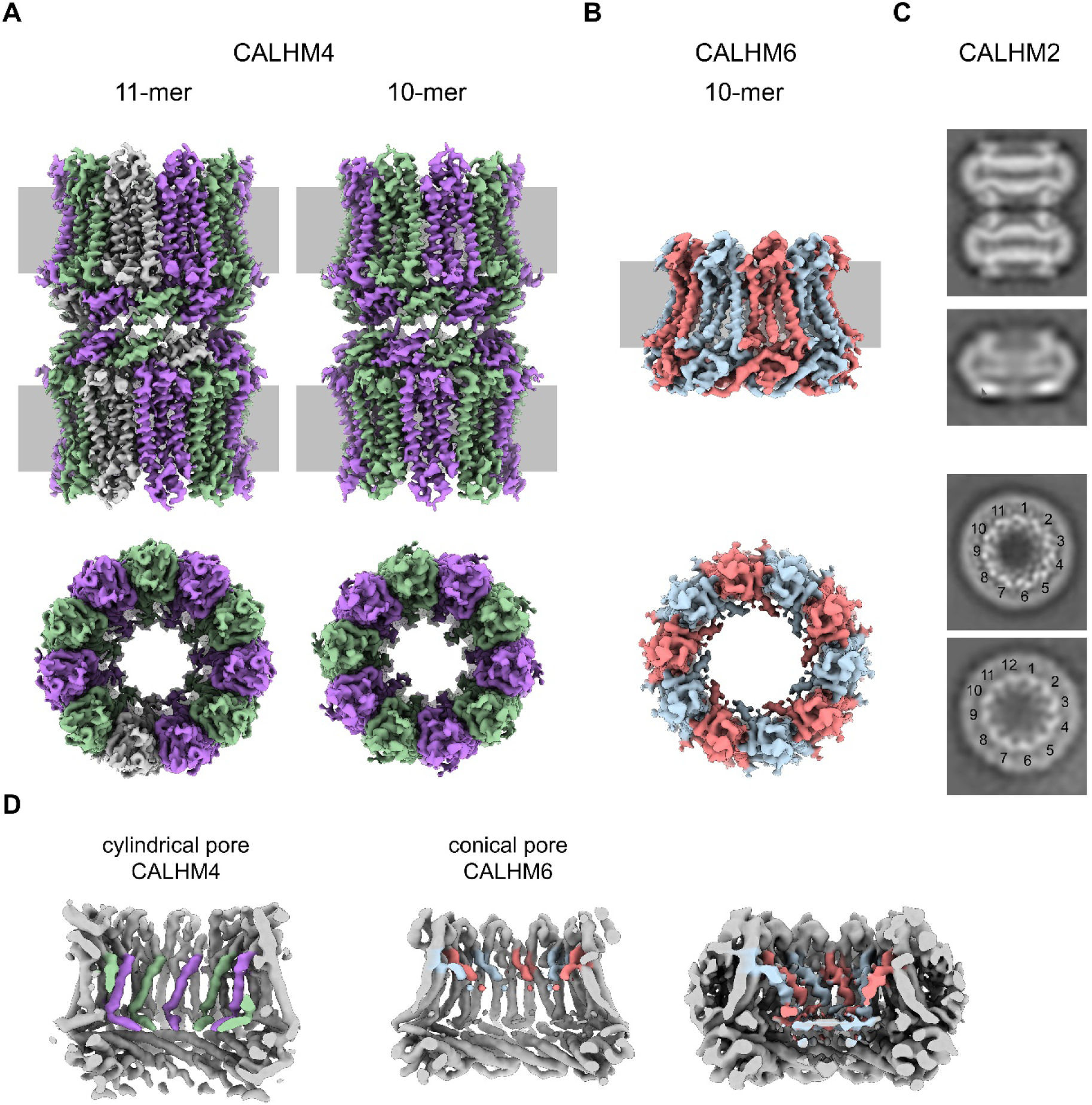
Cryo-EM analysis. (**A**) Cryo-EM density of undecameric (11-mer) and decameric (10-mer) pairs of CALHM4 channels at 3.9 and 4.1 Å respectively. Data was recorded from a sample containing Ca^2+^. Subunits are colored in lilac and green, respectively. (B) Cryo-EM density of decameric CALHM6 channels at 4.5 Å. Subunits are colored in red and light-blue, respectively. A, B, Views are from within the membrane with membrane indicated as grey rectangle (top) and from the outside (bottom). (**C**) Selected 2D classes of the CALHM2 data showing interacting channel pairs and single channels viewed from within the membrane (top) and views of undecameric and dodecameric channels with subunits numbered (bottom). (D) Slices through the CALHM4 (left) and the CALHM6 (right) channels illustrating the distinct features of the cylindrical and conical pore conformations. View of CALHM6 at lower contour (right) shows extended density for the mobile TM1. Maps are low-pass filtered at 6 Å. Density corresponding to TM1 is colored according to A and B.

### Cylindrical pore conformation of the CALHM4 structure

For the determination of the CALHM4 structure, we have recorded cryo-EM data in the presence and absence of Ca^2+^ and observed similar properties, which are not affected by divalent cations (Figure 3-figure supplement 2 and 3). In each case, we found a heterogeneous, about equal distribution of particles with two distinct oligomeric states. The smaller particles are composed of decameric and the larger of undecameric assemblies, both of which we refer to as CALHM4 channels (Figure 4). Due to the slightly higher quality of the Ca^2+^-containing sample, we continued to use this data for further analysis. The cryo-EM density of the decamers extends to a resolution of 4.2 Å and undecamers to 3.9 Å, which in both cases permitted the unambiguous interpretation by an atomic model (Figure 3-figure supplements 2, 3 and 5A, Video 1, Table 1). In each structure, the subunits are arranged around a central axis of symmetry that defines the ion-conduction path. As described before, both assemblies contain pairs of CALHM4 channels of the same size related by two-fold symmetry that interact via their cytoplasmic parts (Figure 4-figure supplement 1A-C). The CALHM4 channels form approximately 90 Å high cylindrical proteins that span the lipid bilayer with regions at their respective periphery extending into the aqueous environment on either side of the membrane (Figure 4). In the plane of the membrane, the dimensions of decameric and undecameric CALHM4 channels amount to 110 Å and 120 Å respectively (Figure 4, Figure 4-figure supplement 1D-E). Irrespective of their distinct oligomerization, the individual subunits in both assemblies show equivalent conformations, which resemble the recently described structures of CALHM1 and 2 (Choi et al., 2019; Demura et al., 2020; Syrjanen et al., 2020) and thus define an organization that is likely general for the CALHM family (Figure 5A, Figure 5-figure supplement 1). Although the members of this family were predicted to share their architecture with connexins and related innexin and LRRC8 channels (Siebert et al., 2013), this turns out not to be the case. Whereas subunits in all families contain four transmembrane helices, which run perpendicular to the membrane, the mutual arrangement of these α-helices differs between CALHM channels and their connexin-like counterparts. When viewed from the extracellular side, the transmembrane helices in connexin, innexin and VRAC channels (Deneka et al., 2018; Maeda et al., 2009; Oshima et al., 2016) are arranged in a clockwise manner, whereas the arrangement in CALHM channels is anticlockwise (Figure 5B, C). Moreover, in connexins and related channels, the four helices form a tightly interacting left-handed bundle with a common core, resulting in a structure with an oval cross-section (Figure 5C). Conversely, the helices of the CALHM subunit are organized as two layers conferring an overall trapezoid cross-section (Figure 5B). Finally, whereas connexins and LRRC8 channels form hexamers, and innexins form octamers, an oligomeric arrangement that has also been observed for CALHM1, the larger CLAHM4 channels contain either 10 or 11 subunits (Figure 4A, B). The outer layer of the CALHM subunit is composed of the interacting α-helices TM2-4, which are arranged in one row sharing mutual interactions only with their respective neighbor. The inner layer consists of TM1, which on its entire length exclusively interacts with TM3, the central helix of the outer layer (Figure 5A, B). When assembled into oligomers, the helices of the outer layer form a ring, which defines the boundaries of the channel (Figure 5B). In this outer ring, the peripheral helices TM2 and 4 are involved in extended interactions with neighboring subunits resulting in a tightly packed interface. Apart from a small fenestration between TM3 and 4 in the center of each subunit, this structural unit shields the interior of the pore from the surrounding membrane (Figure 5-figure supplement 2A). In contrast, the respective TM1 helices forming the inner layer are distant from each other and thus not involved in mutual inter-subunit interactions (Figure 5D, E). In the region preceding TM1, the residues of the N-terminus form a helix (NH) that is oriented perpendicular to the first transmembrane segment parallel to the plane of the lipid bilayer (Figure 4, Figure 5D, Figure 3-figure supplement 5A and Figure 5-figure supplement 2B). Among the family members of known structures, the conformations of NH and TM1 are best defined in CALHM4 (Figure 3-figure supplement 5A), whereas weaker density of these fragments in the equivalent state of CALHM1 and 2 points towards a higher mobility of this region in latter proteins (Choi et al., 2019; Syrjanen et al., 2020). On the extracellular side, a short loop bridges α-helices TM1 and 2 and an extended region containing two short α-helices (E1H and E2H), connects TM3 with the long TM4, which extends beyond the membrane plane (Figure 5A). The extracellular domain is stabilized by two conserved disulfide bridges connecting both extracellular segments at the C-terminal end of TM1 in the observed conformation (Figure 5-figure supplement 2C). On the intracellular side, TM2 precedes a short helix (CLH) that projects away from the pore axis with TM3 being bent in the same direction (Figure 5-figure supplement 2D). Both α-helices are connected by a 12-residue long loop that is probably mobile and thus not defined in the density. Downstream of TM4, we find an extended intracellular region. The first halve of this region consists of a long α-helix (CTH), which is tilted by 70° towards the membrane plane and which tightly interacts with the corresponding parts of neighboring subunits thus forming a 30 Å-high intracellular ring (Figures 4 and 5A, Figure 4-figure supplement 1C). Distal to CTH, a 62-residue long extended loop, which includes partly defined in the cryo-EM density interspersed secondary structures,, folds back towards the intracellular ring on the outer surface of the channel with its C-terminus extending to the opposed subunit of the juxtaposed CALHM4 channel (Figure 4-figure supplements 1B, C). These contacts form the bulk of the interaction relating CALHM4 channel pairs in an arrangement whose relevance in a cellular context is still ambiguous.

**Figure 4.**
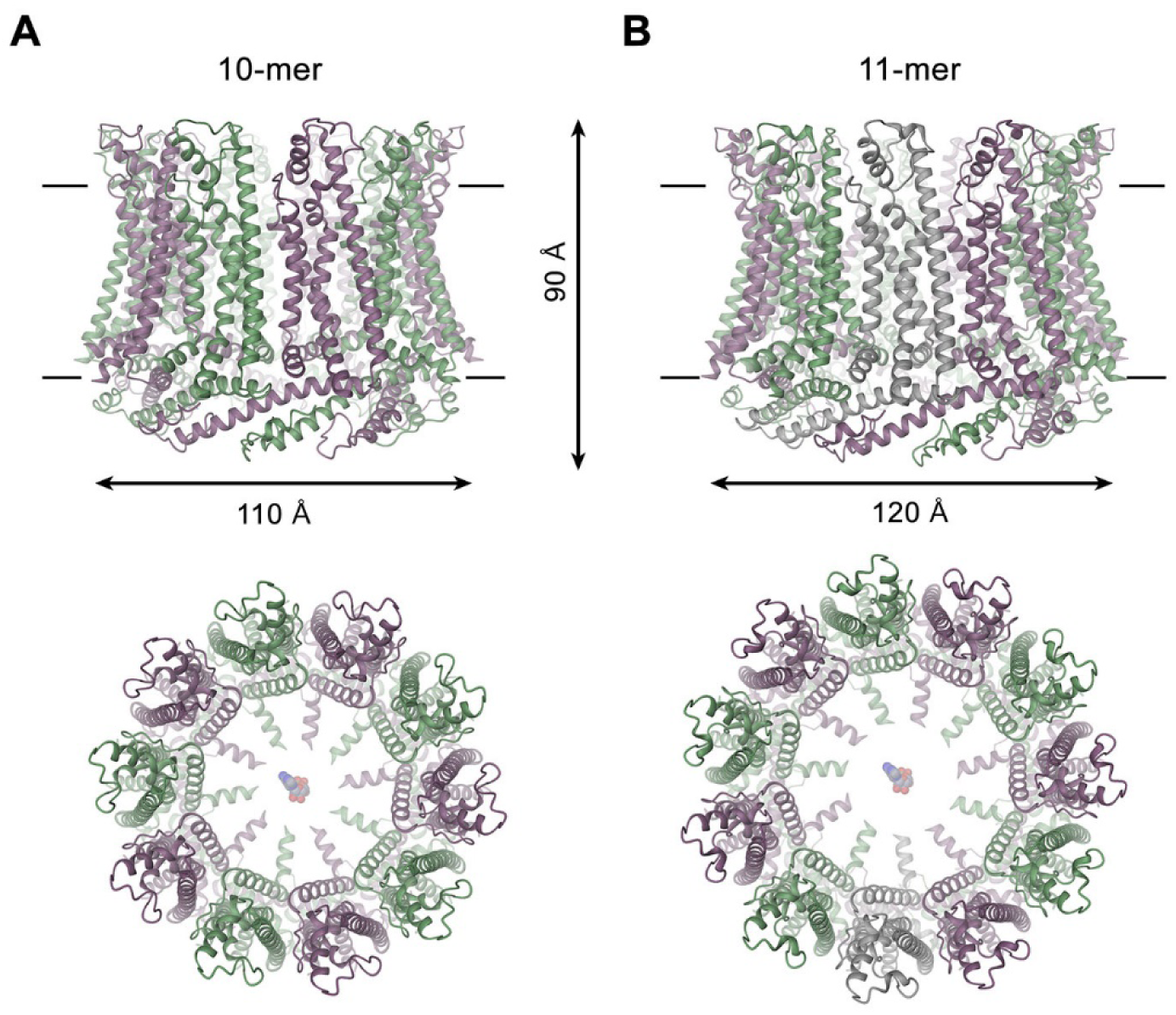
CALHM4 structure. Ribbon representation of (**A**), decameric and (B), undecameric CALHM4 channels. Top views are from within the membrane with membrane boundaries indicated, bottom views are from the extracellular side with an ATP molecule placed for reference. The approximate dimensions are indicated. Subunits are colored in lilac and green.

**Figure 5.**
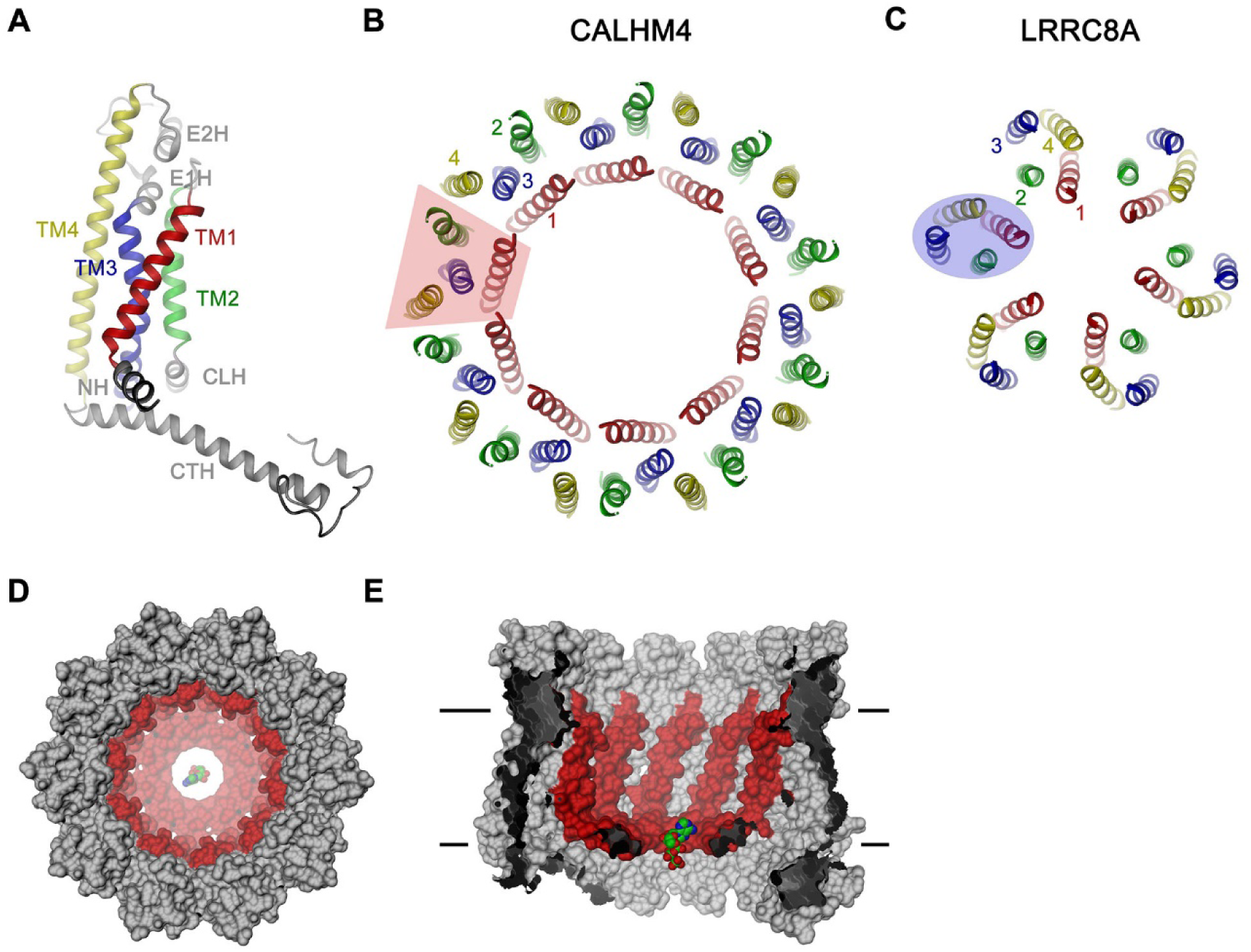
CALHM4 subunit and oligomeric arrangement. (**A**) Ribbon representation of the CALHM4 subunit. Secondary structure elements are labeled and transmembrane α-helices are shown in unique colors. View of the transmembrane α-helices of (**B**), the CALHM4 decamer and (**C**), the volume regulated anion channel LRRC8A from the extracellular side. B-C, color code is as in A, transmembrane segments of one subunit are numbered. The general shape of a single subunit is indicated (B, trapezoid, C, oval). (**D**) Surface representation of the CALHM4 decamer. The view is from the outside. (**E**), Slice through the CALHM4 pore viewed from within the membrane. D, E, TM1 and the N-terminal α-helix NH are colored in red, ATP is shown as a space-filling model for comparison.

**Table 1.**
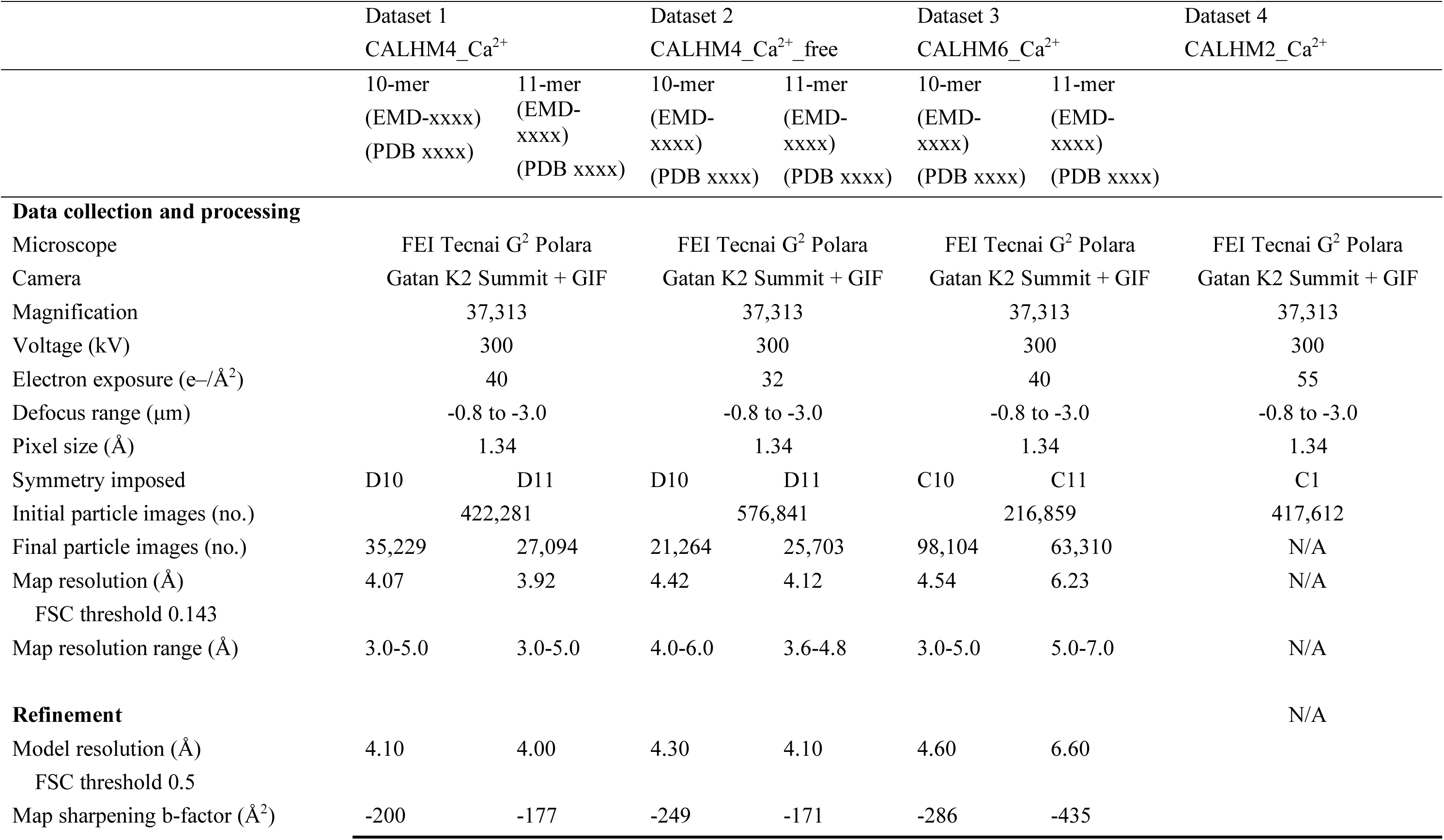
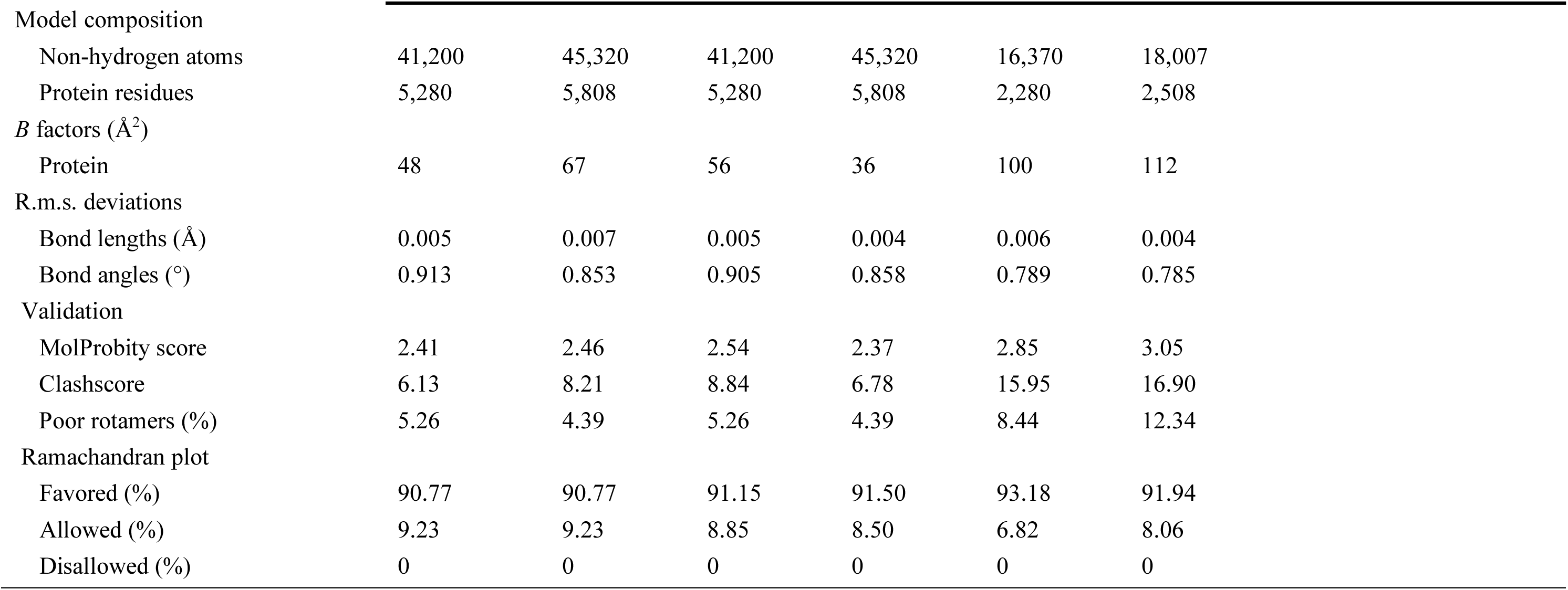
Cryo-EM data collection, refinement and validation statistics.

|  | Dataset 1 |  | Dataset 2 |  | Dataset 3 |  | Dataset 4 |
| --- | --- | --- | --- | --- | --- | --- | --- |
|  | CALHM4_Ca<sup>2+</sup> |  | CALHM4_Ca<sup>2+</sup>_free |  | CALHM6_Ca<sup>2+</sup> |  | CALHM2_Ca<sup>2+</sup> |
|  | 10-mer<br>(EMD-xxxx)<br>(PDB xxxx) | 11-mer<br>(EMD-xxxx)<br>(PDB xxxx) | 10-mer<br>(EMD-xxxx)<br>(PDB xxxx) | 11-mer<br>(EMD-xxxx)<br>(PDB xxxx) | 10-mer<br>(EMD-xxxx)<br>(PDB xxxx) | 11-mer<br>(EMD-xxxx)<br>(PDB xxxx) |  |
| **Data collection and processing** |  |  |  |  |  |  |  |
| Microscope | FEI Tecnai G<sup>2</sup> Polara |  | FEI Tecnai G<sup>2</sup> Polara |  | FEI Tecnai G<sup>2</sup> Polara |  | FEI Tecnai G<sup>2</sup> Polara |
| Camera | Gatan K2 Summit + GIF |  | Gatan K2 Summit + GIF |  | Gatan K2 Summit + GIF |  | Gatan K2 Summit + GIF |
| Magnification | 37,313 |  | 37,313 |  | 37,313 |  | 37,313 |
| Voltage (kV) | 300 |  | 300 |  | 300 |  | 300 |
| Electron exposure (e-/Å<sup>2</sup>) | 40 |  | 32 |  | 40 |  | 55 |
| Defocus range (µm) | -0.8 to -3.0 |  | -0.8 to -3.0 |  | -0.8 to -3.0 |  | -0.8 to -3.0 |
| Pixel size (Å) | 1.34 |  | 1.34 |  | 1.34 |  | 1.34 |
| Symmetry imposed | D10 | D11 | D10 | D11 | C10 | C11 | C1 |
| Initial particle images (no.) | 422,281 |  | 576,841 |  | 216,859 |  | 417,612 |
| Final particle images (no.) | 35,229 | 27,094 | 21,264 | 25,703 | 98,104 | 63,310 | N/A |
| Map resolution (Å) | 4.07 | 3.92 | 4.42 | 4.12 | 4.54 | 6.23 | N/A |
| FSC threshold 0.143 |  |  |  |  |  |  |  |
| Map resolution range (Å) | 3.0-5.0 | 3.0-5.0 | 4.0-6.0 | 3.6-4.8 | 3.0-5.0 | 5.0-7.0 | N/A |
| **Refinement** |  |  |  |  |  |  | N/A |
| Model resolution (Å) | 4.10 | 4.00 | 4.30 | 4.10 | 4.60 | 6.60 |  |
| FSC threshold 0.5 |  |  |  |  |  |  |  |
| Map sharpening b-factor (Å<sup>2</sup>) | -200 | -177 | -249 | -171 | -286 | -435 |  |

|  |  |  |  |  |  |  |
| --- | --- | --- | --- | --- | --- | --- |
| Model composition |  |  |  |  |  |  |
| Non-hydrogen atoms | 41,200 | 45,320 | 41,200 | 45,320 | 16,370 | 18,007 |
| Protein residues | 5,280 | 5,808 | 5,280 | 5,808 | 2,280 | 2,508 |
| *B* factors (Å<sup>2</sup>) |  |  |  |  |  |  |
| Protein | 48 | 67 | 56 | 36 | 100 | 112 |
| R.m.s. deviations |  |  |  |  |  |  |
| Bond lengths (Å) | 0.005 | 0.007 | 0.005 | 0.004 | 0.006 | 0.004 |
| Bond angles (°) | 0.913 | 0.853 | 0.905 | 0.858 | 0.789 | 0.785 |
| Validation |  |  |  |  |  |  |
| MolProbity score | 2.41 | 2.46 | 2.54 | 2.37 | 2.85 | 3.05 |
| Clashscore | 6.13 | 8.21 | 8.84 | 6.78 | 15.95 | 16.90 |
| Poor rotamers (%) | 5.26 | 4.39 | 5.26 | 4.39 | 8.44 | 12.34 |
| Ramachandran plot |  |  |  |  |  |  |
| Favored (%) | 90.77 | 90.77 | 91.15 | 91.50 | 93.18 | 91.94 |
| Allowed (%) | 9.23 | 9.23 | 8.85 | 8.50 | 6.82 | 8.06 |
| Disallowed (%) | 0 | 0 | 0 | 0 | 0 | 0 |

### Pore architecture

Decameric and undecameric CALHM4 channels contain wide pores, which are cylindrical throughout except for a constriction at the intracellular membrane boundary. The diameter at both entrances of the pore measures about 52 Å and 60 Å for decameric and undecameric assemblies respectively, thus defining the properties of an unusually large channel, which could be permeable to molecular substrates (Figure 6A, Figure 4-figure supplement 1D, E). Even at the constriction located at the intracellular membrane leaflet where the respective N-terminal helices NH project towards the pore axis, the decameric channels are about 20 Å and undecameric channels 30 Å wide (Figure 6A). In both cases the pore would thus be sufficiently large to accommodate an ATP molecule, consistent with the notion of CALHM proteins as proteins forming ATP-permeable channels (Figures 4, 5D, E, Figure 4-figure supplement 1D, E). Remarkably, this large pore size is in sharp contrast to our functional characterization by electrophysiology where we did not observe appreciable currents for CALHM4 (Figure 2A, Figure 2-figure supplement 1E), which suggests that either the absence of activating or presence of inhibiting components might impede ion conduction.

**Figure 6.**
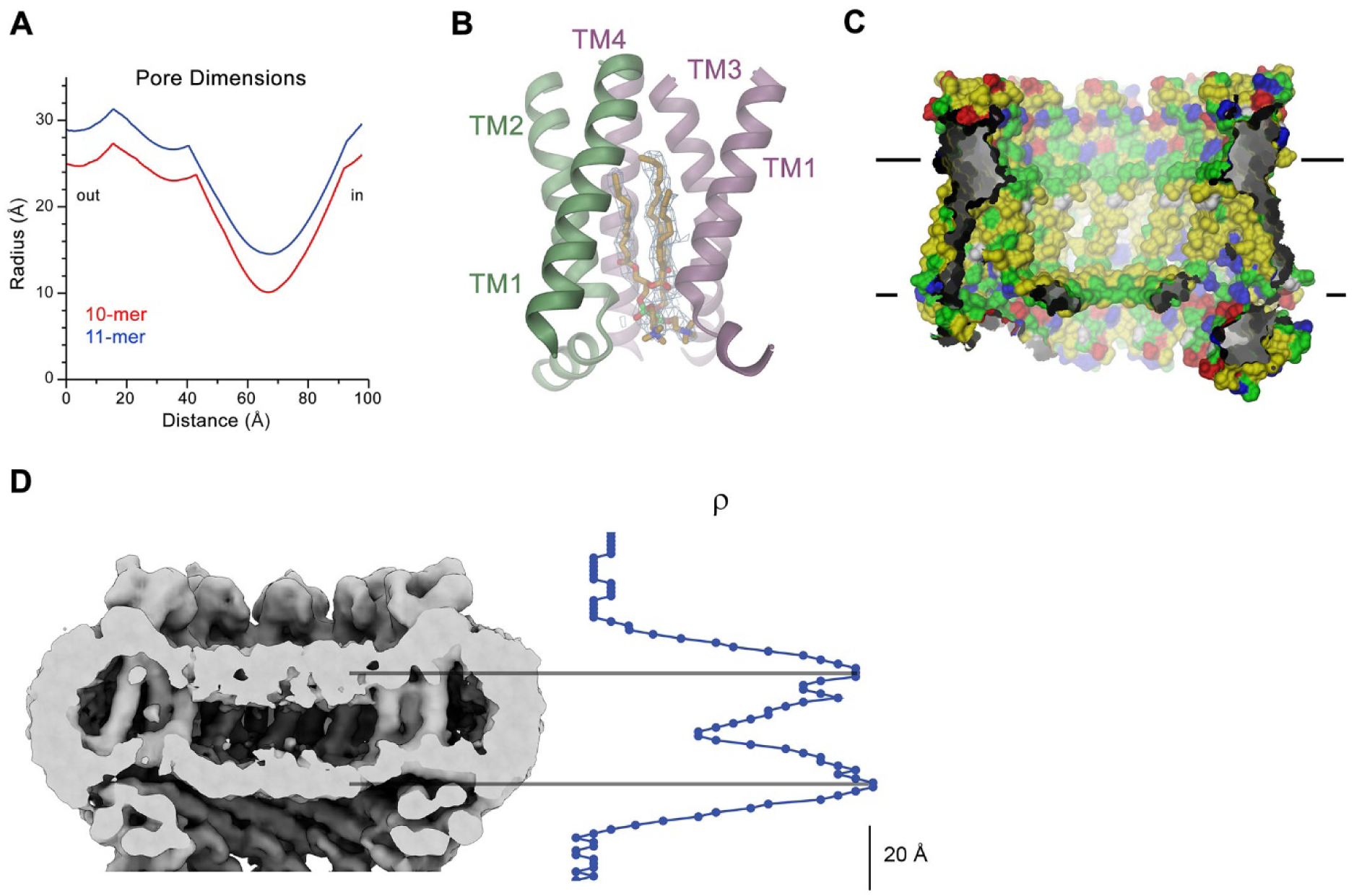
CALHM4 pore properties and lipid interactions. (**A**), Pore radius of the CALHM4 decamer (red) and undecamer (blue) as calculated in HOLE (Smart et al., 1996). (**B**) Two phosphatidylcholine molecules modeled into residual cryo-EM density (blue mesh) in a cavity at the interface between neighboring α-helices TM1. Secondary structure elements are indicated. (**C**) Chemical properties of residues lining the pore of the CALHM4 channel. Shown is a slice through the pore viewed from within the membrane. The protein is displayed as molecular surface. Hydrophobic residues are colored in yellow, polar residues in green, acidic residues in red and basic residues in blue. (**D**) Slice through the pore region of the CALHM4 decamer viewed from within the membrane. Shown is non-averaged density in a single copy of the decameric CALHM4 channel pair at low contour to highlight the location of increased density within the pore corresponding to a bilayer of either phospholipids or detergents. A plot of the density along the pore axis showing two maxima that are separated by the expected distance between the headgroup regions of lipid bilayer is shown right.

Within the membrane, the pore diameter of CALHM4 is defined by the ring of non-interacting TM1 helices, which is placed inside an outer ring of the channel formed by helices TM2-4 (Figure 5B, D). This arrangement creates clefts between neighboring helices of the inner ring which are delimited by helices TM2 and 4 at the respective subunit interfaces (Figure 5B, E). In our sample, this cleft appears to be filled with lipids as indicated by the residual density observed in the cryo-EM maps (Figure 6B, Figure 6-figure supplement 1A, B). Due to the prevalence of aliphatic residues lining the pore at the assumed location of the membrane core, the pore is highly hydrophobic, whereas the regions extending into the aqueous environment contain polar and charged residues (Figure 6C). Similarly, the side of the helical N-terminus, which faces the membrane at its intracellular boundary, is hydrophobic (Figure 5-figure supplement 2B). In light of its large cross-section and high hydrophobicity, it is conceivable that the interior of the pore would accommodate lipids, which potentially could form bilayers that would restrict ion permeation as recently suggested for CALHM2 based on molecular dynamics simulations (Syrjanen et al., 2020). In our data, we find strong evidence for a layered distribution of density inside the pore within the presumed membrane region in non-symmetrized maps and after application of symmetry, which could either reflect the presence of lipids or detergent molecules arranging in a bilayer-like structure facilitated by the confined pore geometry (Figure 6D, Figure 6-figure supplement 1C). The observed density distribution displays features that quantitatively match the corresponding properties of lipid membranes as judged by a comparison to cryo-EM density of liposomes and computer simulations (Figure 6-figure supplement 1D, E). Collectively, our data are compatible with the presence of a membrane located within the pore region of CALHM4 channels, which could interfere with the diffusion of charged substances. Thus, despite of its large pore diameter, it is at this point unclear whether the CALHM4 structure defines a conductive conformation of CALHM channel or alternatively a conformation that facilitates the assembly of a lipid bilayer residing inside the pore that would impede ion conduction.

### Conical pore conformation of the CALHM6 structure

As in case of CALHM4, data of CALHM6 shows a heterogeneous distribution of decameric and undecameric assemblies, but in this case with a prevalence of the former amounting to 60% of the classified particles (Figure 3-figure supplement 4). Both assemblies of CALHM6 contain subunits with equivalent conformations, which are better defined in the smaller oligomers (Figure 3B, Figure 3-figure supplement 4). We have thus chosen the decameric CALHM6 structure for a description of the conical pore of a CALHM channel (Figures 3D and 7A-C). Unlike CALHM4, the lower resolution of the CALHM6 density of 4.5 Å, precludes a detailed interpretation of the structure for all parts of the protein. Still, the high homology between the two paralogs and density attributable to large side chains constrains the placement of helices and conserved loop regions and thus allows the credible analysis of major conformational differences (Figure 3-figure supplement 5B, Video 2, Table 1). Unlike CALHM4, there is no dimerization of CALHM6 channels and the C-terminal region following the cytoplasmic helix CTH instead appears to engage in intramolecular interactions with the outside of the cytoplasmic rim for most of its length (Figure 7A). This observation further supports the notion that the dimerization of CALHM4 might be a consequence of interactions formed between solubilized proteins where the mobile C-terminus could equally well engage in intra- and intermolecular interactions, of which the latter would be multiplied in the highly symmetric arrangement. Although the general organization of the CALHM6 channel closely resembles CALHM4, it shows a distinctly different state of the pore (Figure 3D). Upon comparison of decameric channels of both paralogs, in CALHM6 we find a slight expansion of the protein parallel to the membrane and an accompanying moderate contraction in perpendicular direction (Figure 7-figure supplement 1A, Video 3). In a superposition of the subunits, similar conformations are observed for α-helices TM2 and 4 and the C-terminal helix CTH and larger differences in the interacting helices TM1 and 3 (Figure 7D, Figure 7-figure supplement 1B, C). These differences are most pronounced for TM1, which in CALHM6 has detached from TM3 and moved by 60° towards the pore axis around a hinge located upstream of a conserved phenylalanine at the end of TM1 (Figure 7B-E). In both structures, the conformation of the proximal loop connecting TM1 and 2 is stabilized by two conserved cysteines, which are involved in disulfide bridges with the region connecting TM3 and TM4 (Figure 7F-G, Figure 5-figure supplement 2C). As a consequence of the disruption of its interaction with TM1, the intracellular halve of TM3 tilts away from the pore axis by 30° around a pivot located close to a conserved proline residue (P115 in CALHM4), which probably destabilizes the helix (Figure 7-figure supplement 1B, D). The transition from a conformation observed in CALHM4 to a conformation defined by CALHM6 is accompanied by the dissociation of interactions between TM1 and 3, which are mediated by conserved residues involving a cluster of hydrophobic interactions at the extracellular side and additional interactions on the entire length of both α-helices (Figure 7F-G, Figure 7-figure supplement 1C, D). The iris-like movement of TM1 in the oligomeric channel in its transition between conformations probably requires concerted rearrangements to avoid steric clashes in the crowded environment of the pore. When viewed from the outside this transition thus resembles the closing of an aperture and it converts a cylindrical pore to a funnel which narrows towards the intracellular side while creating a large cavity between TM1 and TM3 that becomes accessible from the cytoplasm (Figure 7C, Video 3). In the cryo-EM density, TM1 is less well defined compared to the rest of the protein reflecting its increased flexibility in the observed structure (Figure 3-figure supplement 5B). Since the conformation of NH, which has moved towards the pore axis, is not defined in the density, the size of the CALHM6 channel at its constriction remains ambiguous. It could range from an occluded pore if NH helices make contacts in the center of the channel (modeled as clogged conformation) to a pore with similar diameter as found at the CALHM4 constriction. The latter could be obtained in case of the mutual relationship between TM1 and NH being unchanged compared to the CALHM4 structure (defined as kinked conformation) or in case of straightening of NH in continuation of TM1 (in an extended conformation) (Figure 7-figure supplement 1E). In any case, the large conformational changes would affect the location of lipids within the pore, which in case of an internal bilayer would have to rearrange in response to the severely altered pore geometry. Such rearrangement is reflected in the changed distribution of the residual electron density inside the pore (Figure 7H). Whereas, compared to CALHM4, we find density at the location of the intracellular layer of lipids and further towards the cytoplasm, part of which might be attributable to the mobile N-terminal α-helix NH, the outer layer of density corresponding to the putative extracellular leaflet of a bilayer has disappeared. Thus, despite the pronounced conformational differences to CALHM4 and the fact that the CALHM6 structure appears to contain features of a closed pore, its definitive functional assignment remains also in this case ambiguous.

**Figure 7.**
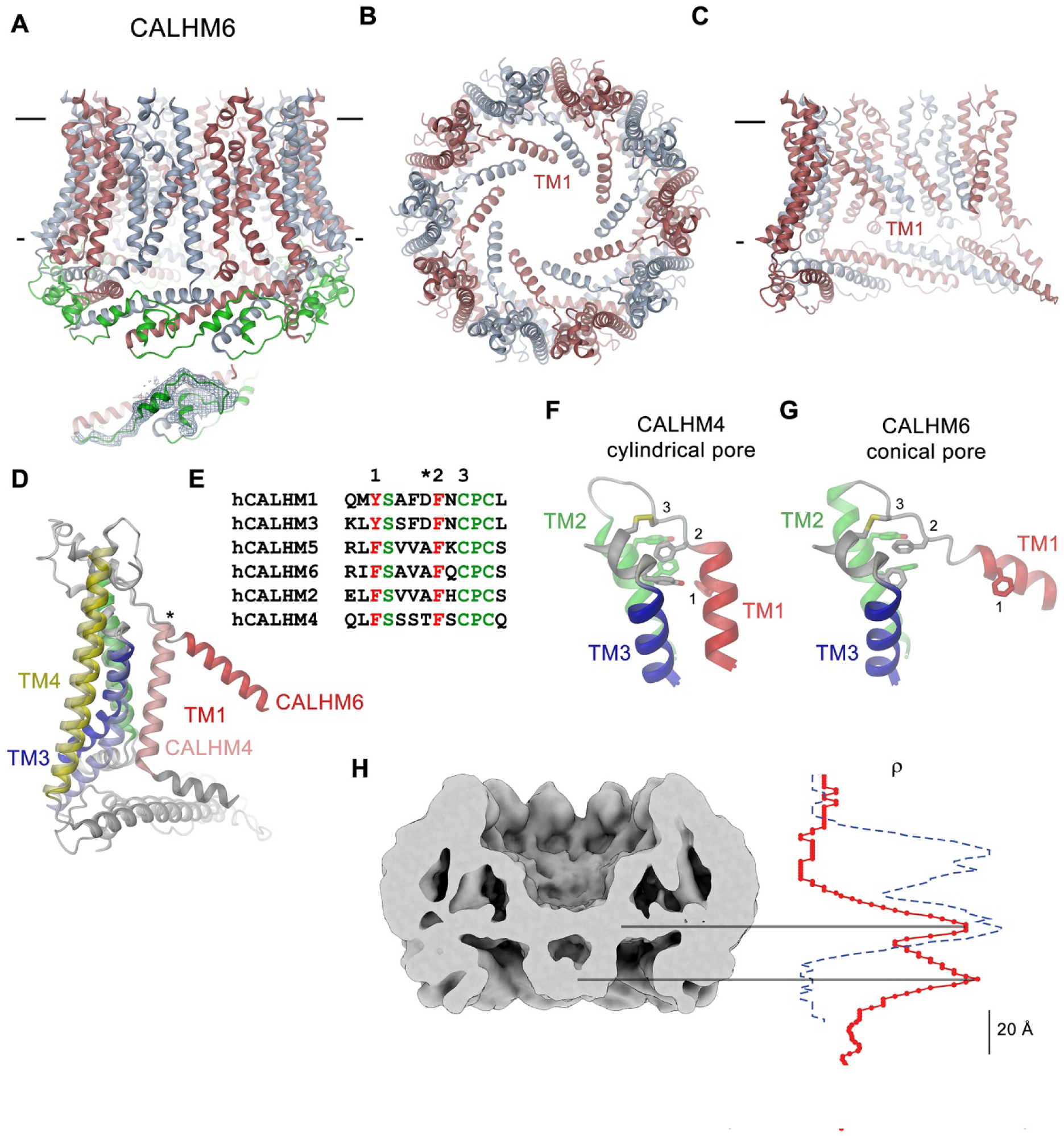
CALHM6 structure. (**A**) Ribbon representation of the decameric CALHM6 structure viewed from within the membrane. Subunits are colored in red and light blue. Part of the mobile C-terminus defined in low-pass filtered cryo-EM density that was modeled as Ala and Gly chain is colored in green. Inset (below) shows a close-up of this region with cryo-EM density superimposed. (**B**) View of the CALHM structure from the extracellular side. (**C**) Slice through the pore of the CALHM6 structure. B, C, TM1, which has moved towards the pore axis is labeled. A, C, membrane boundary is indicated. (**D**) Superposition of single subunits of the CALHM4 and CALHM6 structures illustrating conformational changes. Coloring is as in Figure 5A with CALHM4 shown in brighter shades of the same color. Secondary structure elements are labeled and hinge for the movement of TM1 is indicated by asterisk. (**E**) Sequence alignment of the end of TM1 and the following loop of CALHM paralogs. Selected conserved residues are colored in green and red. Numbering corresponds to residues indicated in panels F and G. Asterisk marks a hinge region displayed in D. Close-up of the extracellular region involved in conformational changes in (**F**), the cylindrical conformation displayed in CALHM4 and (**G**), the conical conformation displayed in CALHM6. Residues contributing to a cluster of aromatic residues on TM1-3 and a conserved disulfide bridge are shown as sticks. Selected positions highlighted in E are labeled. F, G, Coloring is as in Figure 5A. (**H**) Slice through the pore region of the CALHM6 decamer viewed from within the membrane. Shown is non-averaged density at low contour to highlight the location of diffuse density within the pore. A plot of the density along the pore axis of CALHM6 is shown in red, the corresponding density in CALHM4 is shown as a dashed blue line for comparison. The two maxima in the CALHM6 density are shifted towards the intracellular side. The density corresponding to the headgroups of the outer leaflet of the bilayer in CALHM4 is absent. Density at the location of the headgroup region at the inner leaflet of the bilayer and further towards the intracellular side could correspond to either lipids or to the poorly ordered N-terminus.

## Discussion

In the presented study, we have addressed the structural and functional properties of CALHM channels in the human placenta. To identify relevant CALHM paralogs in this organ, we have quantified their expression in samples of the whole placenta and in isolated trophoblast cells and found CALHM2, 4 and 6 to be abundant on a transcript level (Figure 1). Their high expression compared to other membrane proteins (Figure 1-figure supplement 1) suggests an important functional relevance of these membrane channels in mediating transport processes during the development of the fetus, although their detailed role still awaits to be explored.

The structural characterization of the three placental paralogs by cryo-electron microscopy has provided insight into fundamental features of the family which largely conform with properties that have recently been described for CALHM1 and 2 (Choi et al., 2019; Demura et al., 2020; Syrjanen et al., 2020) but which also show substantial differences. Similar to previous studies, our structures have defined the architecture of the CALHM subunit which, although containing the same number of membrane-spanning helices as VRACs and gap-junctions forming connexins, innexins and pannexins, is distinct from these proteins (Figure 5B, C). The CALHM subunits thus constitute unique modular building blocks that assemble with different stoichiometries into large membrane channels. In contrast to previously described CALHM structures where the oligomeric state of individual channels was described as uniform, our data shows heterogeneous populations with CALHM4 and CALHM6 assembling as decameric and undecameric channels with similar abundance and CALHM2 as predominantly undecameric proteins with a smaller population of dodecamers (Figure 3, Figure 3-figure supplements 2-4 and 6). The diverse oligomerization of different paralogs is generally consistent with the assemblies observed for the previously described CALHM1 and CALHM2 structures, which were reported to form octamers and undecamers, respectively (Choi et al., 2019; Demura et al., 2020; Syrjanen et al., 2020), and it underlines the ability of CALHM proteins to constitute membrane channels of different sizes. Although the physiological role of these different oligomeric states is currently unclear, the observed heterogeneity might reflect the low energetic penalty for the incorporation of additional subunits into large oligomeric channels of the CALHM family. In this respect, the smaller size of CALHM1 channels could be responsible for its functional properties, which are manifested in electrophysiological recordings. Whereas CALHM1 in our hands showed the previously described functional hallmarks of channels that are activated by depolarization and removal of extracellular calcium, we have not observed pronounced activity of CALHM2, 4 and 6 under the same conditions, despite their efficient targeting to the plasma membrane (Figure 2, Figure 2-figure supplement 1). The low current response contrasts with the larger oligomeric organization of these proteins, which should lead to channels of even higher conductance than observed for CALHM1, and thus likely reflects their low open probability. Together our findings suggest that CALHM2, 4, and 6 are regulated by distinct, still unknown mechanisms. In that respect it remains puzzling how the large pores observed for the investigated structures (with diameters of CALHM4 decamers exceeding 40 Å within the membrane and 20 Å at the respective constriction, Figure 6A) can be regulated to prevent leakage of substances under resting conditions, which would be deleterious to the cell. Despite the unknown activating stimuli, our study has provided insight into gating transitions of CALHM channels by showing two conformations with either a cylindrical pore of uniform large dimeter within the membrane that is constricted at the intracellular side as in case of CALHM4 and a conical pore that continuously narrows from its extracellular entry towards the cytoplasm as in case of CALHM6 (Figures 4 and 7). The main difference in the pore geometry results from a large rearrangement of the pore-lining α-helix TM1 which is accompanied by a smaller change in TM3 (Figure 7D). Whereas in the cylindrical pore conformation of CALHM4, TM1 tightly interacts with TM3, which is a part of a densely packed outer rim of the pore consisting of TM2-4, the helix has dissociated from its interaction site and instead moved towards the symmetry axis to alter the pore geometry in the conical conformation of CALHM6. Similar conformational properties have previously been described for CALHM2 channels, although with a different orientation of TM1 in the conical pore conformation (Choi et al., 2019) (Figure 8-Figure supplement 1A-C). In this previous study, the cylindrical conformation was assigned to an open state and the conical conformation to a closed state of the pore, a proposal that is further supported by a structure of the bound blocker rubidium red which appears to stabilize the conical conformation (Choi et al., 2019) (Figure 8). While, at first glance, the relationship between observed pore conformations and their corresponding functional states appears evident, there are still puzzling questions which prevent a definitive assignment at this stage. *E.g.*, in light of the large diameter of the CALHM4 pore, the poor conductance properties in functional recordings remain mysterious. In that respect, the presence of lipids within the pore of large CALHM channels is noteworthy as it offers a potential alternative mechanism for regulation. In our structure of CALHM4, we find bound lipids inside the pore to stabilize the gap between non-interacting TM1 helices from different subunits (Figure 6B). Additionally, the bimodal diffuse residual cryo-EM density within the hydrophobic interior of CALHM4, which either originates form detergents or co-purified lipids that assemble as bilayers in the constrained environment of the pore are remarkable (Figure 6D). This distribution of density has changed markedly in the conical structure of CALHM6 (Figure 7H) and hints at a potential role of lipids in shaping the activation and permeation properties of large CALHM channels, which is currently still not understood (Figure 8). We thus propose two alternative models for channel regulation in large CALHM channels where the gate in the closed structure of a CALHM1 channels could either be formed by part of the protein or by lipids (Figure 8, Figure 8-Figure 1D). Another remarkable feature of large CALHM channels concerns their tendency to form pairwise assemblies upon extraction from the membranes (Figure 3A, C, Figure3-figure supplements 2, 3 and 6). This property has previously been observed for undecameric CALHM2 channels, which were described to dimerize on the extracellular side thus forming structures that resemble gap-junctions, and in the present study where dimerization was found to some degree in all samples. In our data, the described behavior is most pronounced in case of CALHM4 where dimerization proceeds on the intracellular side mediated by interactions with the partly mobile C-terminus. Although intriguing, it is currently unclear whether any of the observed oligomerizations is of relevance in a physiological context.

**Figure 8.**
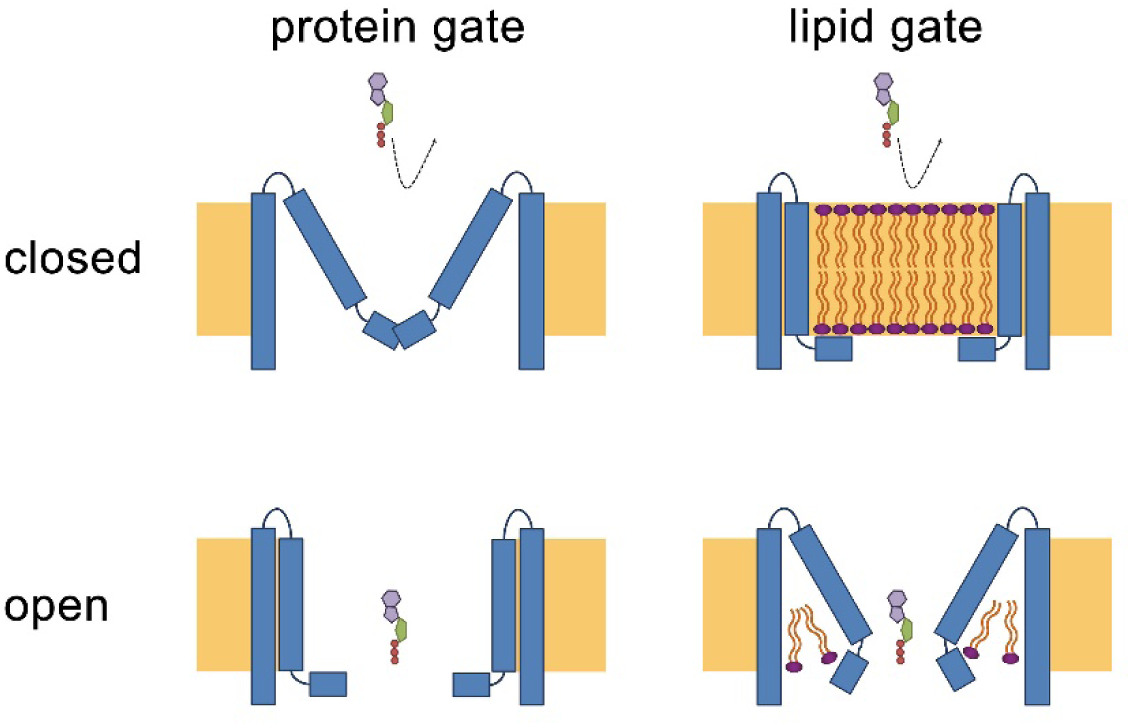
Hypothetical gating mechanisms. Schematic illustration of hypothetical gating mechanisms of large CALHM channels. Left, ‘protein gate’. The gate impeding ion conduction of large CALHM channels (i.e. CALHM2, 4, 6) in the closed state is formed by the N-terminus of the protein, which closes the pore as observed in the modeled ‘clogged’ conformation of CALHM6 (top). In this case the conical conformation of CALHM6 would display a closed pore and the cylindrical conformation (bottom) an open pore. Right, ‘lipid gate’. The gate impeding ion conduction of large CALHM channels in the closed state is formed by lipids assembling as a bilayer within the protein, which impedes ion conduction. Since bilayer formation is facilitated in the cylindrical conformation of CALHM4, this structure represents a closed pore (top) whereas the structure of the bilayer would be disturbed in the conical conformation of CALHM6 (bottom). Since both the ‘kinked’ and ‘extended’ pore conformations of CALHM6 show a large opening, these conformations could represent open pores.

Thus, although our data has provided a large step forward towards the comprehension of CALHM channels, it has also opened many questions. An important area of future investigations relates to the characterization of the localization of CALHM channels in different placental cell types and the identification of their subcellular distribution. Knowledge of their localization would provide a first glimpse into potential roles of these large channels for transport processes in this organ and should provide answers on the potential relevance of intra- and extracellular interactions of channel pairs. Another question concerns the ability of the three paralogs to heteromerize and the potential relevance of such heteromeric channels in a physiological environment. Finally, it will be important to study the activation mechanism of the described channels and whether they are embedded in larger interaction networks, which shape their activation properties. These topics will be addressed in future studies for which our current data provides an important foundation.

## Material and Methods

### Key resources table

| Reagent type<br>(species) or<br>resource | Designation | Source or reference | Identifier |
| --- | --- | --- | --- |
| Antibodies | Mouse Monoclonal Cytokeratin 7 Antibody (OV-TL12/30) [Alexa Fluor® 488] | Bio-Techne | Cat#NBP2-47940AF488 |
| Antibodies | Mouse Monoclonal Vimentin Antibody (V9) [Alexa Fluor® 647] | Bio-Techne | Cat#NBP1-97670AF647 |
| Antibodies | Rabbit anti-CALHM2 | Novusbio | Cat#NBP1-87936 |
| Antibodies | Rabbit anti-CALHM4 | Pineda Antibody-Service | N/A |
| Antibodies | Rabbit anti-CALHM6 | Biorbyt | Cat#orb106933 |
| Antibodies | Goat anti-Rabbit (HRP) | Abcam | Cat#ab6721 |
| Chemical compound, drug | HyClone HyCell TransFx-H medium | GE Healthcare | Cat#SH30939.02 |
| Chemical compound, drug | Phosphate Buffered Saline | Sigma | Cat#806552 |
| Chemical compound, drug | L-glutamine | Sigma | Cat#G7513 |
| Chemical compound, drug | Penicillin-streptomycin | Sigma | Cat#P0781 |
| Chemical compound, drug | Fetal bovine serum | Sigma | Cat#F7524 |
| Chemical compound, drug | Poloxamer 188 solution | Sigma | Cat#P5556 |
| Chemical compound, drug | Polyethylenimine 25K MW, branched | Sigma | Cat#408727 |
| Chemical compound, drug | Polyethylenimine hydrochloride, 40K MW, linear | Polysciences | Cat#24765 |
| Chemical compound, drug | Dulbecco's Modified Eagle's Medium (DMEM) High glucose, pyruvate | Gibco | Cat#41966-029 |
| Chemical compound, drug | Valproic acid sodium salt | Sigma | Cat#P4543 |
| Chemical compound, drug | cOmplete, EDTA-free Protease Inhibitor Cocktail | Roche | Cat# |
| Chemical compound, drug | Glycol-diosgenin (GDN) | Anatrace | Cat#GDN101 |
| Chemical compound, drug | Lauryl Maltose Neopentyl Glycol (LMNG) | Anatrace | Cat#NG310 |
| Chemical compound, drug | D-desthiobiotin | Sigma | Cat#D1411 |
| Chemical compound, drug | Triton X-100 | Millipore Sigma | Cat#T9284 |
| Chemical compound, drug | Biotin | Millipore Sigma | Cat#B4501 |
| Chemical compound, drug | Tween 20 | Sigma | Cat#93773 |
| Commercial assay or kit | GoScript™ Reverse Transcriptase | Promega | Cat#A5003 |
| Commercial assay or kit | GoTaq® qPCR Master Mix for Dye-Based Detection | Promega | Cat#A6002 |
| Commercial assay or kit | NucleoBond PC 10000 | Macherey-Nagel | Cat#740593 |
| Commercial assay or kit | StrepTactin Superflow affinity resin slurry | IBA Lifesciences | Cat#2-1206-002 |
| Commercial assay or kit | Superose 6 10/300 GL | GE Healthcare | Cat#17-5172-01 |
| Commercial assay or kit | Superose 6 5/150 | GE Healthcare | Cat#29091597 |
| Commercial assay or kit | 200 mesh Au 1.2/1.3 cryo-EM grids | Quantifoil | Cat#N1-C14nAu20-01 |
| Commercial assay or kit | 200 mesh Au 0.6/1 cryo-EM grids | Quantifoil | Cat#N1-C11nAu20-01 |
| Commercial assay or kit | Amicon 100 kDa MWCO centrifugal filter | Millipore | Cat#UFC810096 |
| Commercial assay or kit | 0.22 µm Ultrafree-MCCentrifugal Filter | Millipore | Cat#UFC30GV |
| Commercial assay or kit | RNeasy Mini Kit | Qiagen | Cat#74104 |
| Commercial assay or kit | mMESSAGE mMACHINE™ SP6 Transcription Kit | Ambion | Cat#AM1340 |
| Commercial assay or kit | Pierce Cell Surface Protein Isolation Kit | ThermoFisher | Cat#89881 |
| Commercial assay or kit | Amersham ECL Prime Western Blotting Detection Kit | GE Healthcare | RPN2232 |
| Cell line (human) | HEK293S GnTI- | ATCC | CRL-3022 |
| Cell line (human) | HEK-293T | ATCC | CRL-1573 |
| Recombinant DNA | Homo sapiens CALHM1 | GenScript | Accession#NM_001001412.3 |
| Recombinant DNA | Homo sapiens CALHM2 | GenScript | Accession#NM_015916.5 |
| Recombinant DNA | Homo sapiens CALHM3 | GenScript | Accession#NM_001129742.2 |
| Recombinant DNA | Homo sapiens CALHM4 | GenScript | Accession#NM_001366078.1 |
| Recombinant DNA | Homo sapiens CALHM5 | GenScript | Accession#NM_153711.5 |
| Recombinant DNA | Homo sapiens CALHM6 | GenScript | Accession#NM_001010919.3 |
| Recombinant DNA | pTLNX vector | Raimund Dutzler laboratory | Addgene, Cat#49032 |
| Recombinant DNA | Modified pcDNA 3.1 vector with C-terminal 3C protease cleavage site, Venus and Myc tags and streptavidin binding peptide | Raimund Dutzler laboratory | N/A |
| RNA | Antisense Cx38 | Microsynth | (Bahima et al., 2006) |
| Software, algorithm | ViiA™ 7 Software | Applied Biosystems | https://www.thermofisher.com/ch/en/home/life-science/pcr/real-time-pcr/real-time-pcr-instruments/via-7-real-time-pcr-system/via-7-software.html |
| Software, algorithm | SerialEM 3.5.0 | Mastronarde, 2005 | http://bio3d.colorado.edu/SerialEM/ |
| Software, algorithm | RELION-3.0 | Scheres, 2012 | https://www2.mrc-lmb.cam.ac.uk/relion/ |
| Software, algorithm | CTFFIND4.1 | Rohou and Grigorieff, 2015 | http://grigoriefflab.janelia.org/ctf |
| Software, algorithm | Bsoft 1.9.5 | Heymann and Belnap, 2007 | https://lsbr.niams.nih.gov/bsoft/ |
| Software, algorithm | Coot 0.8.8 | Emsley and Cowtan, 2004 | https://www2.mrc-lmb.cam.ac.uk/personal/pemsley/coot/ |
| Software, algorithm | PHENIX 1.14 | Adams et al., 2002 | http://phenix-online.org/ |
| Software, algorithm | REFMAC5 | Murshudov et al., 2011 | http://www.ccpem.ac.uk/ |
| Software, algorithm | MSMS | Sanner et al., 1996 | http://mgltools.scripps.edu/packages/MSMS/ |
| Software, algorithm | DINO 0.9.4 | http://www.dino3d.org | http://www.dino3d.org |
| Software, algorithm | PyMOL 2.3.0 | DeLano, 2002 | https://pymol.org/2/ |
| Software, algorithm | Chimera 1.13.1 | Pettersen et al., 2004 | http://www.cgl.ucsf.edu/chimera/ |
| Software,<br>algorithm | ChimeraX 0.7 | Goddard et al.,<br>2018 | https://www.cgl.ucsf.edu/chimerax/ |
| Software,<br>algorithm | CHARMM | Brooks et al., 1983 | https://www.charmm.org/charmm/ |
| Software,<br>algorithm | Axon Clampex 10.6 | Molecular Devices | N/A |
| Software,<br>algorithm | Axon Clampfit 10.6 | Molecular Devices | N/A |
| Software,<br>algorithm | Prism 7 | GraphPad | https://www.graphpad.com/ |

### Placenta collection

Human placental tissues were collected from the Division of Obstetrics and Gynecology, Lindenhofgruppe Bern, Switzerland, under approval by the ethical commission of the Canton of Bern (approval No Basec 2016-00250). Written informed consent was obtained from all participants. Placentas were collected from uncomplicated pregnancies following elective cesarean section beyond 37 weeks of gestation without prior labor upon patients request or due to breech presentation. All experiments were carried out in accordance with the relevant guidelines and regulations.

### Expression analysis in placental tissue

#### RNA isolation, reverse transcription and quantitative RT-PCR

Approximately 50 mg of frozen placental tissue was subjected to RNA isolation as previously described (Huang et al., 2018; Huang et al., 2013). All RNA samples included in the study had an OD260/280 ratio >1.8. First-strand cDNA was synthesized from 2 μg of total RNA with oligo (dT)_15_ primers and GoScript™ Reverse Transcriptase (Promega, Switzerland) according to the manufacturer’s instructions. The qPCR reaction in SYBR^®^ Green reagent (10 μl) contained 0.5 μM primers, 2 x GoTaq^®^qPCR Master Mix (Promega, Switzerland) and 1 μl cDNA. Primer nucleotide sequences and PCR efficiencies are shown in Figure 1-figure supplement 1C and D. Amplification reactions were performed in duplicates in 384-well plates (Applied Biosystem, USA) with the ViiA7 system (Applied Biosystems, USA). To evaluate mRNA quantities, data were obtained as C_t_ values (describing the cycle number at which logarithmic plots cross calculated threshold lines). C_t_ values were used to determine ΔC_t_ values (ΔC_t_ = C_t_ value of the reference gene minus the C_t_ value of the target gene). The applied reference gene was Tyrosine 3-monooxygenase/tryptophan 5-monooxygenase activation protein, zeta polypeptide (YWHAZ). Comparative transcript data between undifferentiated (CTB) and differentiated (STB) cells were calculated as 2^−ΔΔCt^ values (ΔΔC_t_ = C_t_ value of CALHM gene – C_t_ value of YWHAZ) and are presented as x-fold difference. Data analysis and statistical evaluations were performed using paired 2-way ANOVA with Sidak’s multiple comparisons test with GraphPad Prism® software.

### Primary trophoblast isolation and characterization

Villous trophoblast cells were isolated from healthy human term placentas by enzymatic digestion and gravitational separation as previously described (Nikitina et al., 2011) (Huang et al., 2013) with minor modifications. Briefly, villi-rich tissues were digested three times with 0.25% trypsin (Sigma, USA) and 300 IU/ml Deoxyribonuclease I (Sigma, USA), and then subjected to Percoll^®^ (Sigma, USA) density gradient centrifugation to isolate villous cytotrophoblast cells. In order to assure the purity of the cells, flow cytometry analysis was performed by using the trophoblast-specific epithelial cell marker cytokeratin 7 (Maldonado-Estrada et al., 2004) (anti-cytokeratin 7, Dako, Switzerland). Vimentin (anti-vimentin, Sigma, USA), which is only expressed in potentially contaminating cells (e.g. mesenchymal cells, fibroblasts, smooth muscle cells, stromal cells) served as a negative marker (Soares and Hunt, 2006).

### Primary trophoblast cell culture

Isolated human trophoblast cells were cultured in Dulbecco’s modified Eagle’s medium containing 4.5 g/l glucose (DMEM-HG, Gibco, UK) supplemented with 10% fetal bovine serum (FBS, Seraglob, Switzerland) and antibiotic-antimycotic (Gibco, USA) in a humidified incubator under a 5% CO_2_ atmosphere at 37°C. Cells were seeded at a density of 0.2 10^6^ cells/cm^2^ in costar CellBIND 6 well plates (Corning, USA) and harvested after 24 h (cytotrophoblast stage) or 48 h (STB stage). The syncytialization process was confirmed by visualization under the microscope.

### Expression Cell lines

Adherent HEK293T cells used for protein expression and screening experiments were cultured in DMEM medium supplemented with 10% FBS and penicillin/streptomycin in a humidified incubator under a 5% CO_2_ atmosphere at 37 °C. Suspension-adapted HEK293S GnTI^−^ cells used for protein expression and purification were grown in HyClone TransFx-H media, supplemented with 2% fetal bovine serum (FBS), 4 mM L-glutamine, 1.5% Poloxamer 188 and penicillin/streptomycin in humidified incubator under 5% CO_2_ atmosphere at 37 °C. Both cell lines were tested negative for mycoplasma contamination.

### Construct generation

cDNAs of human CALHM proteins, with SapI restriction sites removed, were obtained from GenScript. For expression in *X. laevis* oocytes, cDNAs of CALHM1, CALHM2, CALHM4 and CALHM6 were cloned into pTLNX vector. For expression in mammalian cells, cDNAs of all human CALHM homologs were cloned into a pcDNA 3.1 vector which was modified to be compatible with FX cloning technology (Geertsma and Dutzler, 2011) and to encode a C-terminal C3 protease cleavage site followed by Venus-Myc-SBP tag.

### Homolog screening

For overexpression studies of CALHM paralogs, adherent HEK293T cells were seeded on 10 cm dishes and grown to a density of 7ˑ10^6^ cells/well. Subsequently, cells were transiently transfected with mixtures of respective *CALHM* cDNA constructs and polyethylenimine with an average molecular weight of 25 kDa (PEI 25K, branched). As a preparative step, 12 µg of DNA and 48 µg of PEI were separately incubated in 0.5 ml of DMEM medium. After 5 min, both solutions were mixed, incubated for 15 min and added to respective cell cultures. 36-h post transfection cells were washed with PBS, harvested by centrifugation at 400 g for 5 min, flash-frozen in liquid nitrogen and stored at −20 °C. For protein extraction, cells were thawed on ice, lysed with lysis buffer (25 mM HEPES, 150 mM NaCl, 0.5 mM CaCl_2_, 2 mM MgCl_2_, 2% GDN, protease inhibitors, RNase, DNase, pH 7.6) for 1 h and clarified by centrifugation at 16,000 g for 10 min. The obtained supernatants were filtered through 0.22 µm centrifugal filters and injected onto a Superose 6 5/150 equilibrated with elution buffer (10 mM HEPES, 150 mM NaCl, 50 µM GDN pH 7.6) and eluted in the same buffer. Proteins were identified by recording of the fluorescence of the attached Venus YFP.

### Protein expression and purification

Suspension HEK293S GnTI^−^ cells were grown in Bioreactor 600 vessels (TPP) and seeded to a density of 0.6 10^6^ cells/ml a day prior transfection. For protein expression, cells were transiently transfected with mixtures of *CALHM* cDNA constructs and PEI MAX 40K. For 300 ml of cell culture, 0.5 mg of DNA and 1.2 mg of PEI MAX were suspended in 20 ml volume of DMEM medium. After 5 min the DNA solutions were mixed with the PEI MAX solutions, incubated for 15 min and supplemented with 4 mM valproic acid. Transfection mixtures were subsequently added to cell cultures and expression was proceeded for 36 h. Afterwards, cells were harvested by centrifugation at 500 g for 15 min., washed with PBS and stored at −20 °C for further use. All following purification steps were carried out at 4 °C. For protein extraction, cells were suspended in lysis buffer (25 mM HEPES, 150 mM NaCl, 1% Lauryl Maltose Neopentyl Glycol (LMNG), 0.5 mM CaCl_2_, 2 mM MgCl_2_, protease inhibitors, RNase, DNase, pH 7.6) and incubated for 1h under constant stirring. For CALHM4 purification in the Ca^2+-^-free conditions, Ca^2+^ was replaced by 5 mM EGTA after extraction. To clarify the extracts, the lysates were centrifugated at 16,000 g for 15 min. Supernatants filtered through 0.5 µm filter were applied to the StrepTactin Superflow affinity resin and incubated with the slurry for 2.5 h under gentle agitation. Unbound proteins were removed and bound proteins were eluted with elution buffer (10 mM HEPES, 150 mM NaCl, 50 µM GDN, 2 mM CaCl_2_ and 10 mM d-Desthiobiotin, pH 7.6). For purification in Ca^2+^-free conditions, Ca^2+^ was replaced by 2 mM EGTA. For cleavage of fusion tags, eluates were incubated for 30 min with C3 protease. Subsequently, samples were concentrated, filtered through 0.22 µm filters and subjected to size exclusion chromatography on a Superose 6 10/300 GL column equilibrated with SEC buffer (10 mM HEPES, 150 mM NaCl, 50 µM GDN, 2 mM CaCl_2_, 2 mM EGTA, pH 7.6). Peak fractions were pooled and concentrated with 100 kDa MWCO centrifugal filters.

### Cryo-EM sample preparation and data collection

For structure determination of human CALHM2, 4 and 6 in the presence of Ca^2+^ and of human CALHM4 in the absence of Ca^2+^ by cryo-EM, 2.5 μl samples of GDN-purified proteins at a concentration of 1.5-3 mg ml^−1^ were applied to glow-discharged holey carbon grids (Quantifoil R1.2/1.3 or R0.6/1 Au 200 mesh). Excess liquid was removed in a controlled environment (4°C and 100% relative humidity) by blotting grids for 4–6 s. Grids were subsequently flash frozen in a liquid propane-ethane mix using a Vitrobot Mark IV (Thermo Fisher Scientific). All samples were imaged in a 300 kV Tecnai G^2^ Polara (FEI) with a 100 μm objective aperture. All data were collected using a post-column quantum energy filter (Gatan) with a 20 eV slit and a K2 Summit direct detector (Gatan) operating in a counting mode. Dose-fractionated micrographs were recorded in an automated manner using SerialEM (Mastronarde, 2005) with a defocus range of –0.8 to –3.0 μm. All datasets were recorded at a nominal magnification of 37,313 corresponding to a pixel size of 1.34 Å/pixel with a total exposure time of 12 s (30 individual frames) and a dose of approximately 1.3 e^−^/Å^2^/frame. The total electron dose on the specimen level for all datasets was approximately between 32 e^−^/Å^2^ and 55 e^−^/Å^2^.

### Cryo-EM image processing

Micrographs from all four datasets were pre-processed in the same manner. Briefly, all individual frames were used for correction of the beam-induced movement using a dose-weighting scheme in RELION’s own implementation of the MotionCor2 algorithm available in version 3.0 (Zivanov et al., 2018). The CTF parameters were estimated on summed movie frames using CTFFIND4.1 (Rohou and Grigorieff, 2015). Low-quality micrographs showing a significant drift, ice contamination or poor CTF estimates were discarded resulting in datasets of 1,125 images of CALHM4 in the presence of Ca^2+^ (dataset 1), 717 images of Ca^2+^-free CALHM4 (dataset 2), 1,059 images of CALHM6 (dataset 3) and 2,065 images of CALHM2 (dataset 4), which were subjected to further data processing in RELION (Scheres, 2012). Particles of CALHM4 and CALHM6 were initially picked using the Laplacian-of-Gaussian method and subjected to 2D classification. 2D class averages showing protein features were subsequently used as templates for more accurate auto-picking as well as input for generating an initial 3D model. From dataset one, 422,281 particles were extracted with a box size of 234 pixels, down-scaled three times and subjected to 2D classification. Having discarded false positives and particles of poor quality, the dataset was reduced to 201,782 particles. Two rounds of non-symmetrized 3D classification using the 60-Å low-pass filtered initial 3D model as a reference allowed to isolate two populations of homogenous particles representing dihedrally-related decameric and undecameric assemblies. These two subsets were refined separately with either D10 or D11 symmetry imposed followed by unbinning to an original pixel size and iterative rounds of 3D refinement, per-particle CTF correction and Bayesian polishing (Zivanov et al., 2018; Zivanov et al., 2019). Extra 3D classification without angular alignment showed increased flexibility within the dimerization interface of interacting intracellular regions in the decameric assembly while no such flexibility was observed for the undecameric population. In order to improve the resolution of the reconstructions, partial signal subtraction was performed, where the density corresponding to the detergent belt as well as the dihedrally-related monomer was subtracted. *In silico* modified particles were auto-refined in the presence of a soft mask around the protein density with either C10 or C11 symmetry imposed. The final map of the decameric assembly at 4.07 Å and of the undecameric assembly at 3.92 Å was sharpened using an isotropic b-factor of –200 Å^2^ and –177 Å^2^, respectively. Dataset of Ca^2+^-free CALHM4 and Ca^2+^-CALHM6 were processed in a similar manner. Briefly, from dataset two, 576,841 particles were extracted and cleaned by 2D classification. The pool, reduced to 97,978 particles, was subjected to non-symmetrized 3D classification that also yielded two populations of dihedrally-related decameric and undecameric assemblies. The final map of the decameric assembly at 4.42 Å and of the undecameric assembly at 4.12 Å was sharpened using an isotropic b-factor of –249 Å^2^ and –171 Å^2^, respectively. From dataset three, 216,859 particles were extracted with a box size of 200 pixels and reduced to 201,761 particles after 2D classification. Non-symmetrized 3D classification also revealed decameric and undecameric populations, although no dihedrally-symmetrized dimers, as in case of CALHM4 in the presence and absence of Ca^2+^, were observed. The final auto-refined map of the decameric assembly at 4.54 Å and of the undecameric assembly at 6.23 Å was sharpened using an isotropic b-factor of – 286 Å^2^ and –435 Å^2^, respectively. In all cases, resolution was estimated in the presence of a soft solvent mask and based on the gold standard Fourier Shell Correlation (FSC) 0.143 criterion (Chen et al., 2013; Rosenthal and Henderson, 2003; Scheres, 2012; Scheres and Chen, 2012). High-resolution noise substitution was applied to correct FSC curves for the effect of soft masking in real space (Chen et al., 2013). The local resolution was estimated using BlocRes from the Bsoft package (Cardone et al., 2013; Heymann and Belnap, 2007).

Visual inspection of micrographs in dataset 4 during pre-processing hinted at preferential orientation of particles. 2D class averages generated from the particles picked with the Laplacian-of-Gaussian method showed primarily views from the extracellular side with a small percentage of side or tilted views, similar to those found in CALHM4 datasets (dihedrally-related dimers) and the CALHM6 dataset (monomers). In order to recover projections at orientations other than from the extracellular side, representative 2D class averages of CALHM4 and 6 were combined and used as 2D templates for auto-picking. The pool of 417,612 particles was subjected to two rounds of 2D classification, after which the dataset was reduced to 71,555 particles. 2D class averages showed that the majority of the remaining particles comprised a view from the extracellular side with only a small fraction representing other orientations. As a consequence of this preferential orientation, projections from other angles were underpopulated and did not show high-resolution features. In other to separate monomeric and dimeric populations, non-symmetrized 3D classification was performed using low-pass filtered reconstructions of CALHM4 and CALHM6. However, both monomeric and dimeric 3D reconstructions of CALHM2 suffered from missing views and therefore were not able to converge to a reliable model during 3D auto-refinement. Additional 2D classification on particles classified either as CALHM2 monomers or dimers showed significant amount of remaining heterogeneity in form of undecameric and dodecameric assemblies, which together with the preferential orientation impeded obtaining a high-resolution reconstruction from this dataset.

### Model building and Refinement

The models of CALHM4 and CALHM6 were built in Coot (Emsley and Cowtan, 2004). CALHM4 was built *de novo* into the cryo-EM density of the undecameric channel and CALHM6 was built using the CALHM4 structure as reference. The cryo-EM density of CALHM4 was of sufficiently high resolution to unambiguously assign residues 4–82 and 94–278. The cryo-EM density of CALHM6 allowed us to assign residues 18–79, 96–129, 146–157 and 162–241. The C-terminal fragment between residues 242–281 was modelled as Gly and Ala chain. The atomic models were improved iteratively by cycles of real-space refinement in PHENIX (Adams et al., 2002) with secondary structure and 22-fold (for CALHM4) and 10-fold (for CALHM6) NCS constraints applied followed by manual corrections in Coot. Validation of the models was performed in PHENIX. Surfaces were calculated with MSMS (Sanner et al., 1996). Figures and videos containing molecular structures and densities were prepared with DINO (http://www.dino3d.org), PyMOL (DeLano, 2002), Chimera (Pettersen et al., 2004) and ChimeraX (Goddard et al., 2018).

### Analysis of the density inside the pore

In order to measure the presence of the density inside the pore, non-symmetrized final 3D reconstructions of CALHM4 and CALHM6 were opened as a stack in Fiji (Schindelin et al., 2012), where the mean density of the area inside the pore was measured along the slices of each 3D reconstruction. The size of the measured area was decided based on the 3D mask generated from the atomic models of CALHM4 and 6. In both cases, the area of 6 x 6 pixels located at the center of the pore axis did not contain any density of the 3D mask meaning that any measured density corresponded to the density outside the mask and therefore did not include any density of CALHM4 or 6.

The ‘experimental’ reference profile was generated based on the electron density map of the MPEG-1 protein bound to a lipidic vesicle (EMD-20622) (Pang et al., 2019). The electron density corresponding to the protein was selected and subtracted in Chimera using the structure of MPEG-1 (PDBID 6U2W). Due to the membrane deformation, the central region of the bilayer was excluded from the measurement. The ‘simulation’ reference profile was generated using the atomistic model of a membrane obtained from MemProtMD database (PDBID 2N5S) (Mineev et al., 2015; Newport et al., 2019). All atoms corresponding to protein, ions and water molecules were removed from the model and the remaining lipids were used to generate an electron density map with 1.34 Å pixel spacing at 6 Å resolution using Chimera (Pettersen et al., 2004). Electron density profiles were generated by measuring a mean pixel intensity of selected regions on map slices along the Z-axis using the Fiji software (Schindelin et al., 2012).

### Two- electrode voltage-clamp recording

For preparation of cRNA coding for CALHM paralogs, *CALHM*-pTLNX DNA constructs were linearized with the FastDigest MluI restriction enzyme (ThermoFisher) purified, transcribed *in vitro* with the mMessage mMachine kit (Ambion) and purified with the RNeasy kit (Qiagen). The obtained cRNAs were either used immediately or aliquoted and stored at 20 °C. Defolliculated *X. laevis* oocytes obtained from Ecocyte Bioscience, were injected with mixtures containing either 1 ng of *CALHM1* cRNA or 5 ng of *CALHM 2, 4 and 6* cRNA and 10 ng of *Xenopus laevis* connexin-38 antisense oligonucleotide (Cx38 ASO) to inhibit endogenous Cx38 currents. Oocytes used as negative control (neg.) were injected with 10 ng of Cx38 ASO only. For protein expression, oocytes were kept in ND69 solution (96 mM NaCl, 2 mM KCl, 1.8 mM CaCl_2_, 1 mM MgCl_2_, 2.5 mM Na-pyruvate, 5 mM HEPES and 50 µg/ml gentamicin, pH 7.5) at 16 °C. Two-electrode voltage clamp (TEVC) measurements were performed 48-60 hours after RNA injections at 20°C. TEVC data were recorded on an OC-725B amplifier (Warner Instrument Corp.) Data recorded at 5 kHz and filtered at 1 kHz were and digitized using a Digidata interface board (1322A or 1440A, Axon Instruments) and analyzed with pCLAMP 10.3 software (Molecular Devices, Sunnyvale, CA). Microelectrode with a resistance of 1–4 MΩ were filled with 3 M KCl. A VC-8 valve controller (Warner Instruments) was used for perfusion of different Ca^2+^ concentrations. Prior to recording, oocytes were additionally injected with a 50 nl of a mixture containing 20 mM BAPTA and 10 mM Ca^2+^solution to minimize activation of Ca^2+^-activated Cl^−^ currents. High Ca^2+^ bath solutions contained, 100 mM Na^+^, 5.4 mM K^+^, 95 mM Cl^−^, 1 mM Mg^2+^, 3 mM Ca^2+^ and 10 mM HEPES, pH 7.2. Divalent cation-free solutions contained 0.5 mM EGTA and 0.5 mM EDTA instead of divalent cations. Intermediate Ca^2+^ concentrations were prepared from both stocks by mixing solution according to the volume calculated with WEBMAXC calculator.

### Cell surface biotinylation

Surface biotinylation of proteins expressed in *X. laevis* oocytes was performed using the Pierce Cell Surface Protein Isolation kit. Oocytes (20-50), injected with *CALHM* cRNAs and incubated for 48-60 hours, were washed 3 times with ND96 solution, transferred to a white 6-well plate (NUNC) and biotinylated by 30 min incubation in 4 ml ND96 solution supplemented with 0.5 mg ml^−1^ EZ-link sulfo-NHS-SS biotin. After incubation, the biotinylation reaction was stopped by addition of quenching solution and oocytes were washed several times with ND96 solution to remove residual reagents. For protein extraction, oocytes were incubated in lysis buffer (25 mM HEPES, 150 mM NaCl, 0.5 mM CaCl_2_, 1% LMNG, 2mM MgCl_2_, protease inhibitors, RNase, DNase, pH 7.6) for 1 h at 4 °C and centrifuged at 10,000 g for 15 min. The supernatants were collected and incubated with NeutrAvidin agarose slurry for 1 h at room temperature under constant mixing. After this step, agarose beads binding the biotinylated proteins were washed with wash solution (10 mM HEPES, 150 mM NaCl, 0.0058% GDN, 2 mM CaCl_2_, protease inhibitors, RNase, DNase). Subsequently, the unbound material was discarded and biotinylated proteins were incubated with SDS-Page sample buffer containing 50 mM DTT for 1 h at RT. Eluted protein fraction was obtained by centrifugation at 1,000 g for 2 min at RT. Membrane protein samples were stored at −20°C. For Western blot analysis, samples were loaded on a 4–20% SDS-polyacrylamide gel. After electrophoretic separation, the proteins were transferred to a polyvinylidene fluoride membrane by a semi-dry blotting procedure. The membranes were first blocked at room temperature for 2 h with 5% non-fat milk in TBS-T buffer (50 mM Tris, 150 mM NaCl, 0.075% Tween20, pH 7.5) and then incubated with respective anti-CALHM primary antibodies overnight at 4°C. To remove unbound primary antibodies, the membranes were washed with TBS-T buffer and subsequently blotted with goat anti-rabbit-HRP conjugated secondary antibody for 2 h at 4 °C. The membranes were washed again with TBS-T buffer and chemiluminescent signals were developed with the Amersham ECL Prime Western Blotting Detection kit.

### Statistics and reproducibility

Paired 2-way ANOVA with Sidak’s multiple comparisons test was applied to detect differences in placental mRNA levels of CALHM isoforms between undifferentiated cytotrophoblast and differentiated syncytiotrophoblast cells. A *p*-value < 0.05 was considered as statistically significant. Statistical comparisons were performed using GraphPad Prism (GraphPad). Electrophysiology data were repeated multiple times with different batches of cRNA and *X. laevis* oocytes with very similar results. Conclusions of experiments were not changed upon inclusion of further data. In all cases, leaky oocytes were discarded. For Ca^2+^ concentration-response analysis using TEVC methods, statistical significance was determined by analysis of variance by Student’s t test. A *p*-value < 0.05 was considered statistically significant. The number of independent experimental repetitions is represented by *n*.

## Acknowledgements

This research was supported by a grant from the Swiss National Science Foundation (No. 31003A_163421) to R.D. We thank O. Medalia and M. Eibauer, the Center for Microscopy and Image Analysis (ZMB) of the University of Zurich, and the Mäxi foundation for the support and access to the electron microscopes, S. Klauser and S. Rast for their help in establishing the computer infrastructure. We are grateful to Rona Karahoda and Saira Shahnawaz for their expert help with qPCR and immunoblotting. The authors want to express their gratitude to the patients, physicians and midwives of the Lindenhofgruppe Bern, who participated in this study. Special thanks also to Dr. Ruedi Moser, Lindenhofgruppe Bern, for coordinating the placenta sampling process. This project was supported by the Stiftung Lindenhof Bern. All members of the Dutzler lab are acknowledged for their help at various stages of the project.

## Accession codes

The cryo-EM density maps of CALHM4 and CALHM6 in presence of Ca^2+^ as well as CALHM4 in absence of Ca2+ have been deposited in the Electron Microscopy Data Bank under ID codes EMD-xxxx, EMD-xxxx and EMD-xxxx, respectively. The coordinates for the atomic model of CALHM4 refined against the 3.92 Å cryo-EM density and the coordinates of CALHM6 refined against the 4.54 Å cryo-EM density have been deposited in the Protein Data Bank under ID codes YYYY and ZZZZ.

## Author Contributions

K.D. generated expression constructs, purified proteins and assisted with structure determination and functional experiments. M.S. prepared the samples for cryo-EM, collected EM data and proceeded with structure determination. M.I.B.S. recorded and analyzed electrophysiology data. D. D. generated and characterized initial constructs. J.Z. and C.A. carried out expression analysis in the placenta. K.D., M. S., M.I.B.S., C.A., J.Z. and R.D. jointly planned experiments, analyzed the data and wrote the manuscript.

## Author Information

The authors declare no competing financial interests. Correspondence and requests for materials should be addressed to R.D. or M.S.

---

**Figure 1—figure supplement 1.**
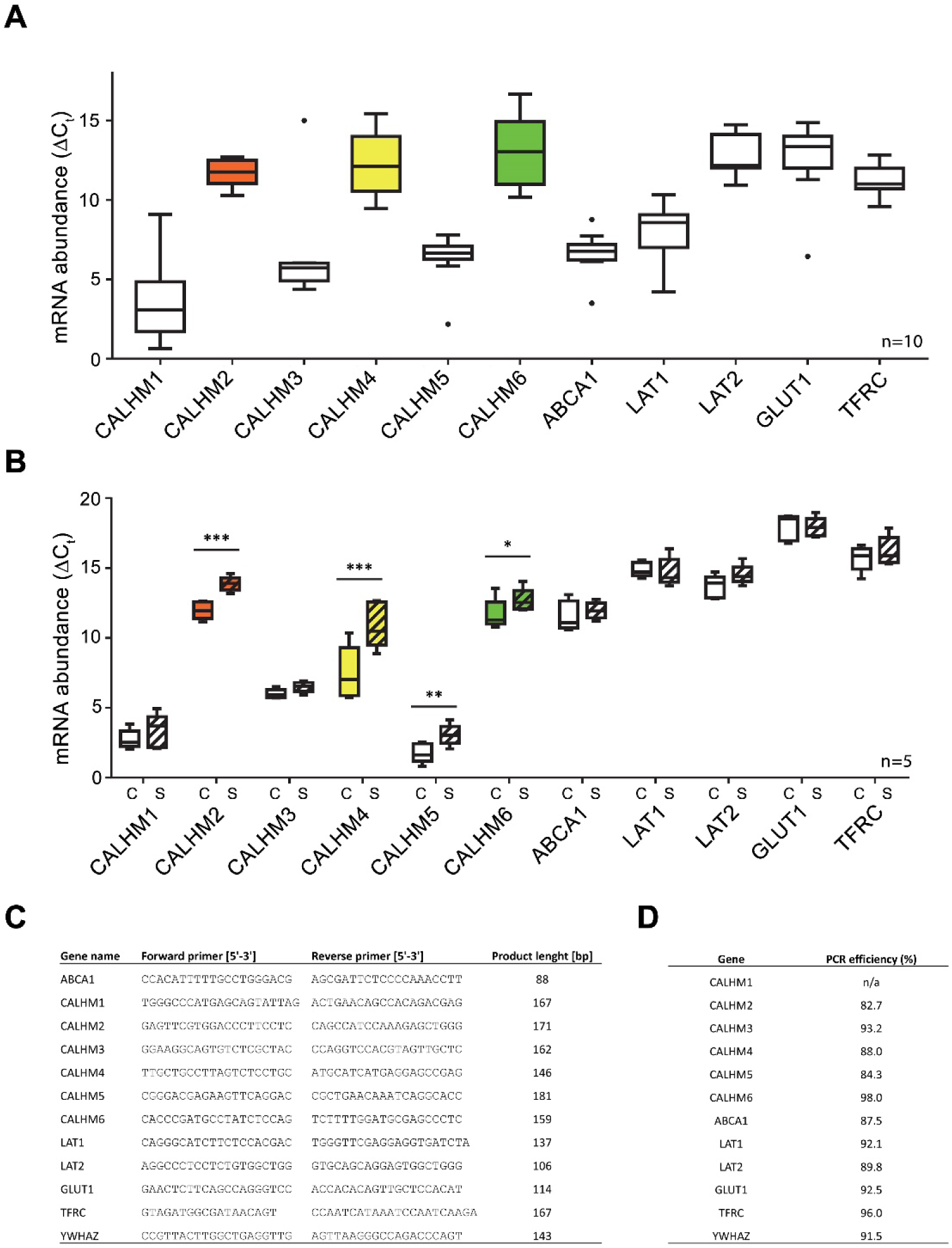
Comparative mRNA expression data for other physiologically relevant transporters/receptors in the human placenta. Y-axis and statistical analysis are according to Figure 1. mRNA concentrations of different CALMH paralogs and of other transport proteins in (**A**), human placental tissues (n=10) and (**B**), human trophoblasts isolated from healthy term placentae (n=5) was assessed by quantitative RT PCR and normalized to the reference gene YWHAZ. Abbreviations: ABCA1, ATP-binding cassette transporter A1; CALHM, calcium homeostasis modulator; LAT1 (SLC7A5), L-type amino acid transporter 1; LAT2 (SLC7A8), L-type amino acid transporter 2; GLUT1 (SLC2A1), Glucose transporter 1; TFRC, transferrin receptor complex 1; YWHAZ, Tyrosine 3-monooxygenase/tryptophan 5-monooxygenase activation protein, zeta polypeptide. (**C**) Primer sequences used for PCR amplification and (**D**), Calculated amplification efficiencies for the indicated genes.

**Figure 2—figure supplement 1.**
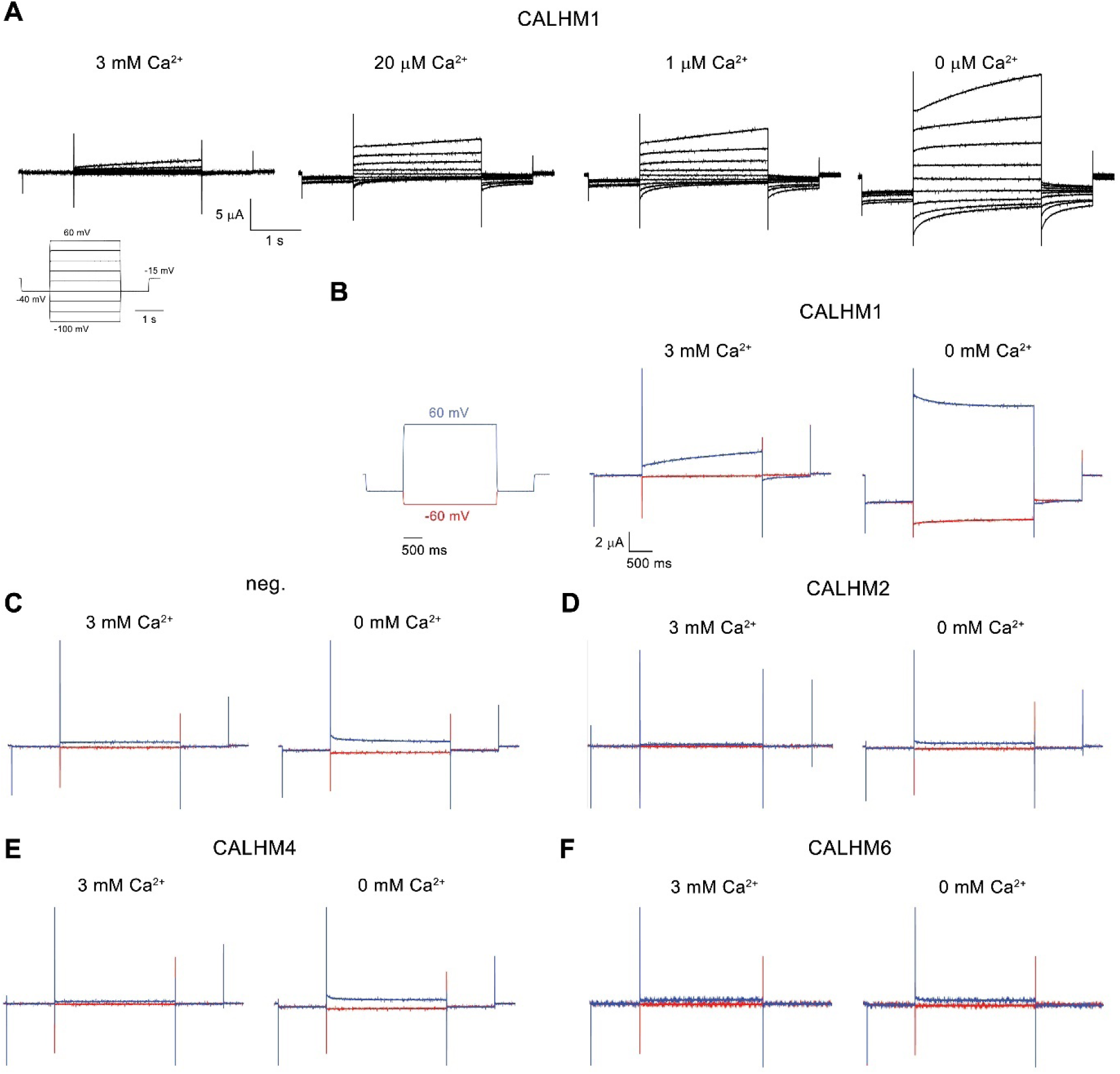
Electrophysiology traces. (**A**) Representative current traces of CALHM1 recorded at indicated extracellular Ca^2+^ concentrations. The voltage protocol is shown as inset (left). (**B-F**) Representative currents of CALHM paralogs (**B**), CALHM1, (**D),** CALHM2, (**E**), CALHM4 and (**F**), CALHM6 in comparison to (**C**), control oocytes (neg.). The current protocol is shown left. Traces including a step to −60 mV are colored in red, traces including a step to 60 mV in blue. A-F, Data were recorded 40-60 hours after injection of cRNA.

**Figure 3—figure supplement 1.**
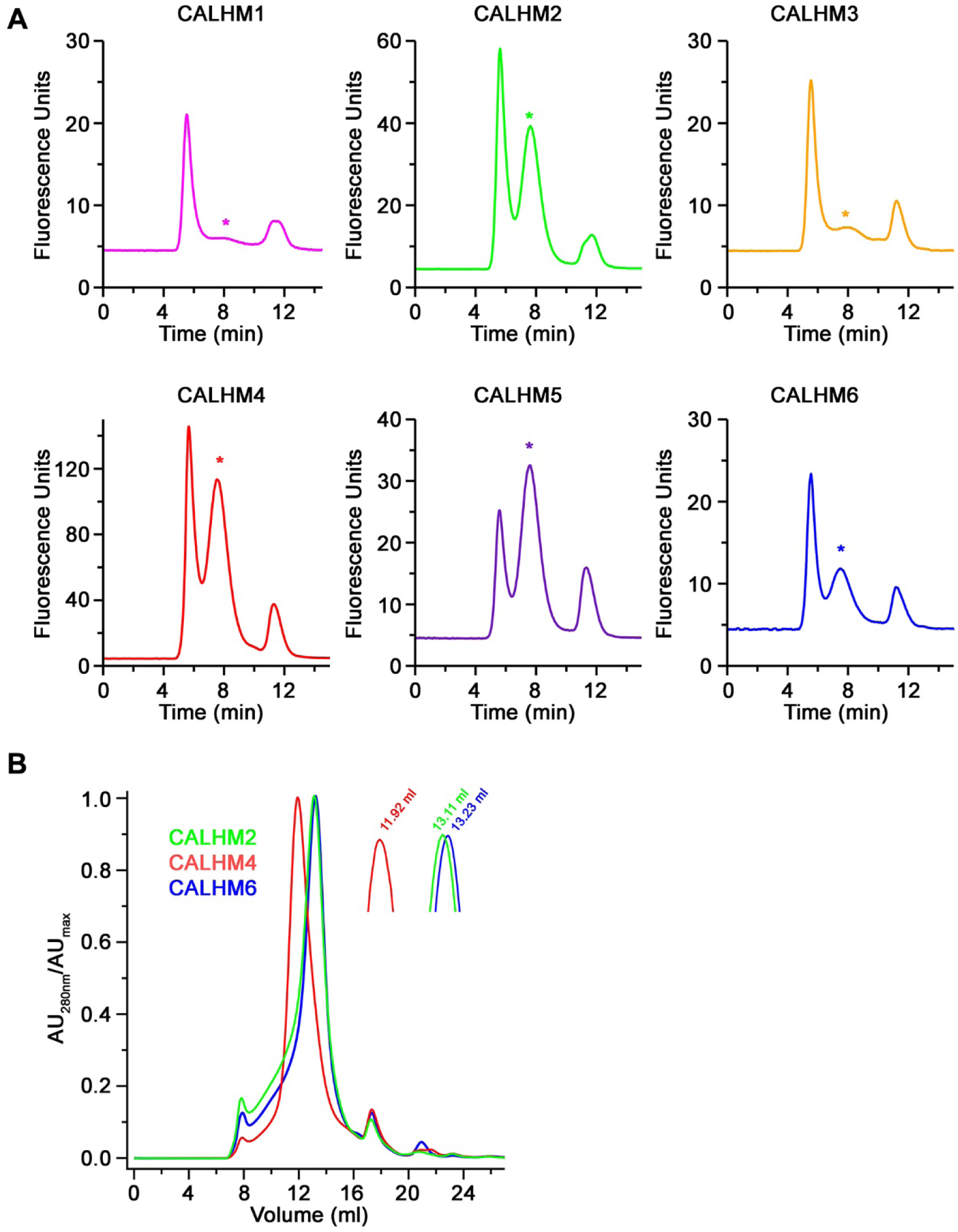
Biochemical characterization. (**A**) Size-exclusion chromatograms recorded from fluorescently tagged CALHM subunits from a whole-cell extract in the detergent GDN on a Superose 6 5/150 column at a flow rate of 0.2 ml/min. Asterisks indicate elution times for each paralog. (**B**) Overlay of size-exclusion chromatograms recorded from purified CALHM paralogs after cleavage of the fusion peptide used for cryo-EM analysis on a Superose 6 10/300 column. Inset shows magnified peak region with respective elution volumes indicated.

**Figure 3—figure supplement 2.**
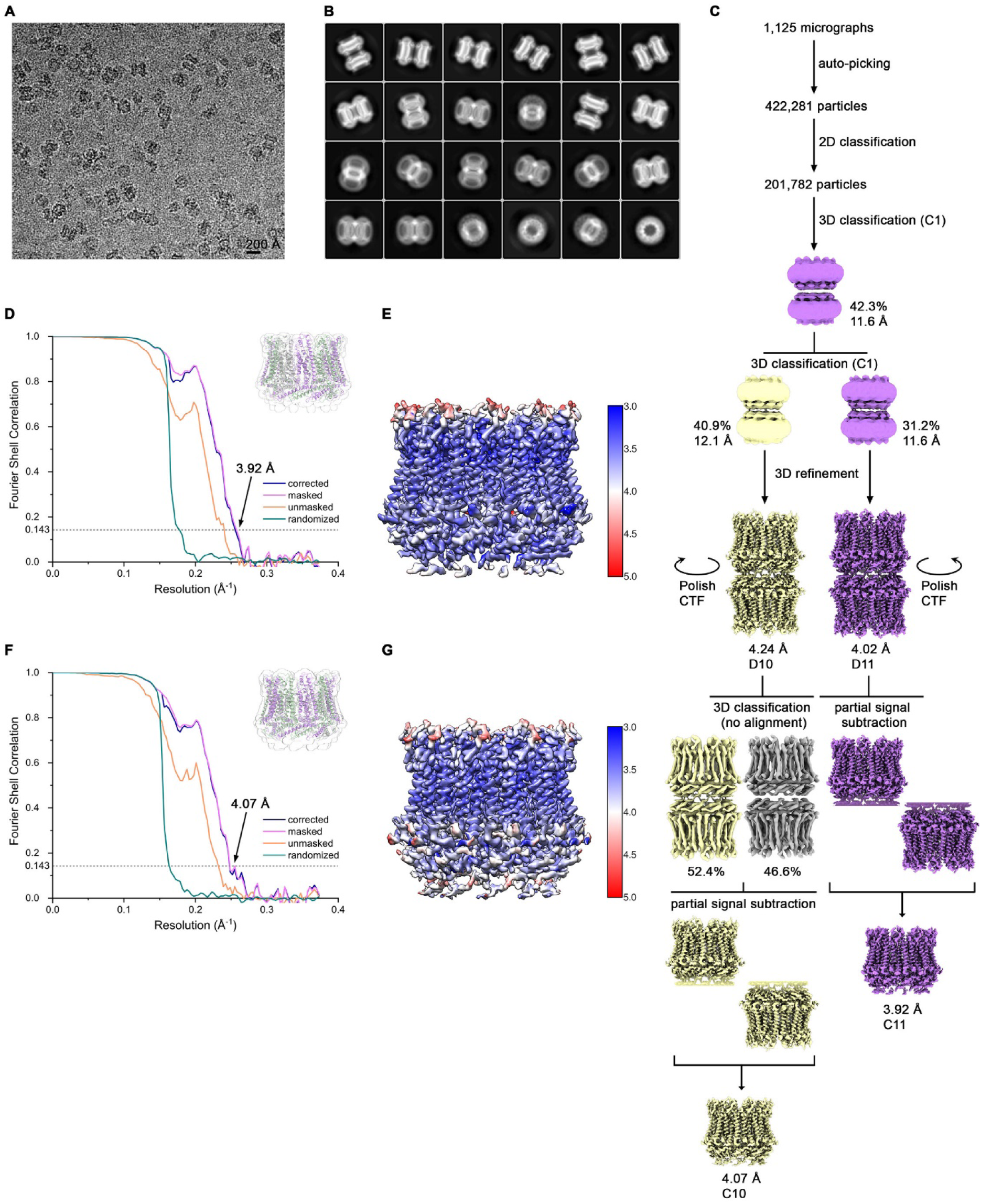
Cryo-EM reconstruction of CALHM4 in presence of Ca^2+^. (**A**) Representative cryo-EM micrograph acquired with a Tecnai G^2^ Polara microscope. (**B**) 2D class averages of CALHM4 in presence of Ca^2+^. (**C**) Data processing workflow. Two rounds of non-symmetrized 3D classification allowed to isolate two populations representing decameric and undecameric assemblies. The particles were further refined with either D10 or D11 symmetry imposed. After performing per-particle CTF refinement and Bayesian polishing, the remaining structural heterogeneity in decamers was segregated by performing the final 3D classification, where the orientations were kept fixed as in the consensus model. To further improve the resolution of each reconstruction, partial signal subtraction followed by 3D refinement was applied. Particles with the symmetry relaxed to either C10 or C11 were merged and subjected to a final round of auto-refinement. The distribution of all particles (%) and the resolution of each class is indicated. (**D**) FSC plot of the final refined undecameric unmasked (orange), masked (pink), phase-randomized (green) and corrected for mask convolution effects (blue) cryo-EM density map of CALHM4. The resolution at which the FSC curve drops below the 0.143 threshold is indicated. The inset shows the atomic model within the mask that was applied for calculations of the resolution estimates. (**E**) Final 3D reconstruction of undecameric CALHM4 colored according to local resolution. (**F**) FSC plot of the final refined decameric unmasked (orange), masked (pink), phase-randomized (green) and corrected for mask convolution effects (blue) cryo-EM density map of CALHM4. The resolution at which the FSC curve drops below the 0.143 threshold is indicated. The inset shows the atomic model within the mask that was applied for calculations of the resolution estimates. (**G**) Final 3D reconstruction of decameric CALHM4 colored according to local resolution.

**Figure 3—figure supplement 3.**
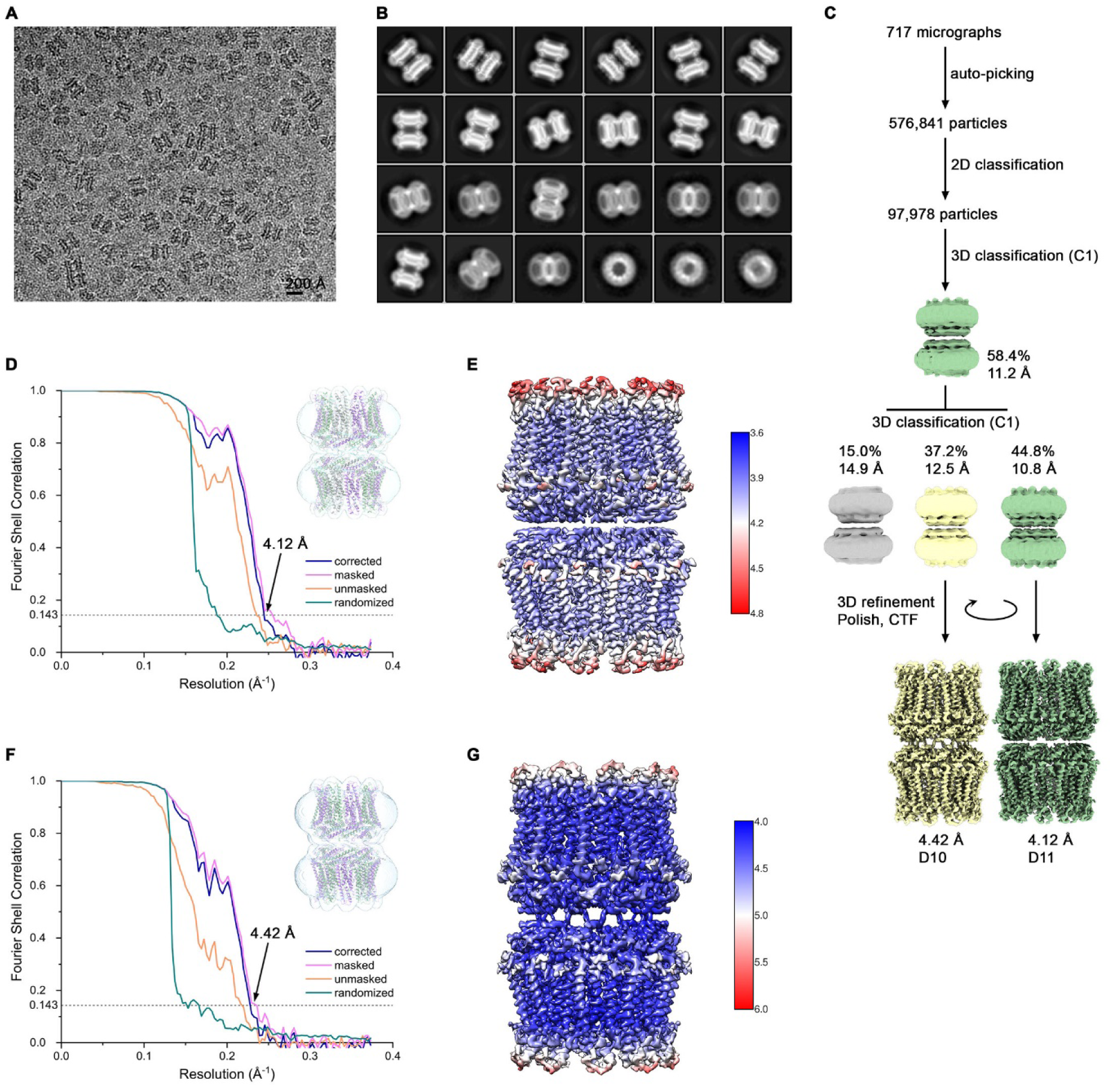
Cryo-EM reconstruction of CALHM4 in absence of Ca^2+^. (**A**) Representative cryo-EM micrograph acquired with Tecnai G^2^ Polara microscope. (**B**) 2D class averages of CALHM4 in absence of Ca^2+^. (**C**) Data processing workflow. Two rounds of non-symmetrized 3D classification allowed to isolate two populations representing decameric and undecameric assemblies. The particles were further refined with either D10 or D11 symmetry imposed and iterative per-particle CTF refinement and Bayesian polishing. The distribution of all particles (%) and the resolution of each class is indicated. (**D**) FSC plot of the final refined undecameric unmasked (orange), masked (pink), phase-randomized (green) and corrected for mask convolution effects (blue) cryo-EM density map of CALHM4. The resolution at which the FSC curve drops below the 0.143 threshold is indicated. The inset shows the atomic model within the mask that was applied for calculations of the resolution estimates. (**E**) Final 3D reconstruction of undecameric CALHM4 colored according to local resolution. (**F**) FSC plot of the final refined decameric unmasked (orange), masked (pink), phase-randomized (green) and corrected for mask convolution effects (blue) cryo-EM density map of CALHM4. The resolution at which the FSC curve drops below the 0.143 threshold is indicated. The inset shows the atomic model within the mask that was applied for calculations of the resolution estimates. (**G**) Final 3D reconstruction of decameric CALHM4 colored according to local resolution.

**Figure 3—figure supplement 4.**
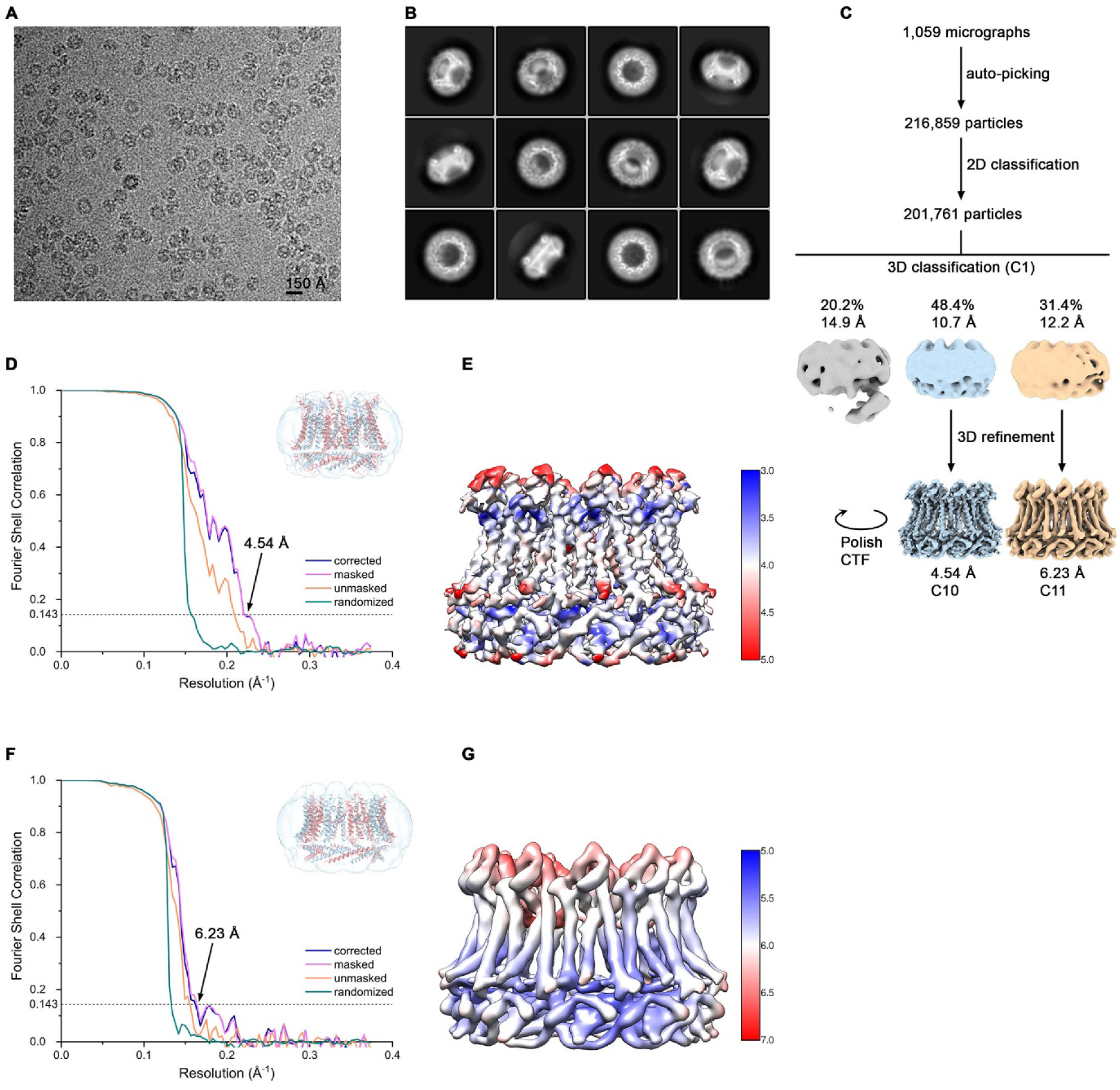
Cryo-EM reconstruction of CALHM6 in presence of Ca^2+^. (**A**) Representative cryo-EM micrograph acquired with a Tecnai G^2^ Polara microscope. (**B**) 2D class averages of CALHM6 in presence of Ca^2+^. (**C**) Data processing workflow. Non-symmetrized 3D classification allowed to isolate two populations representing decameric and undecameric assemblies. The particles were further refined with either C10 or C11 symmetry imposed and iterative per-particle CTF refinement and Bayesian polishing. The distribution of all particles (%) and the resolution of each class is indicated. (**D**) FSC plot of the final refined decameric unmasked (orange), masked (pink), phase-randomized (green) and corrected for mask convolution effects (blue) cryo-EM density map of CALHM6. The resolution at which the FSC curve drops below the 0.143 threshold is indicated. The inset shows the atomic model within the mask that was applied for calculations of the resolution estimates. (**E**) Final 3D reconstruction of decameric CALHM6 colored according to local resolution. (**F**) FSC plot of the final refined undecameric unmasked (orange), masked (pink), phase-randomized (green) and corrected for mask convolution effects (blue) cryo-EM density map of CALHM6. The resolution at which the FSC curve drops below the 0.143 threshold is indicated. The inset shows the atomic model within the mask that was applied for calculations of the resolution estimates. (**G**) Final 3D reconstruction of undecameric CALHM6 colored according to local resolution.

**Figure 3—figure supplement 5.**
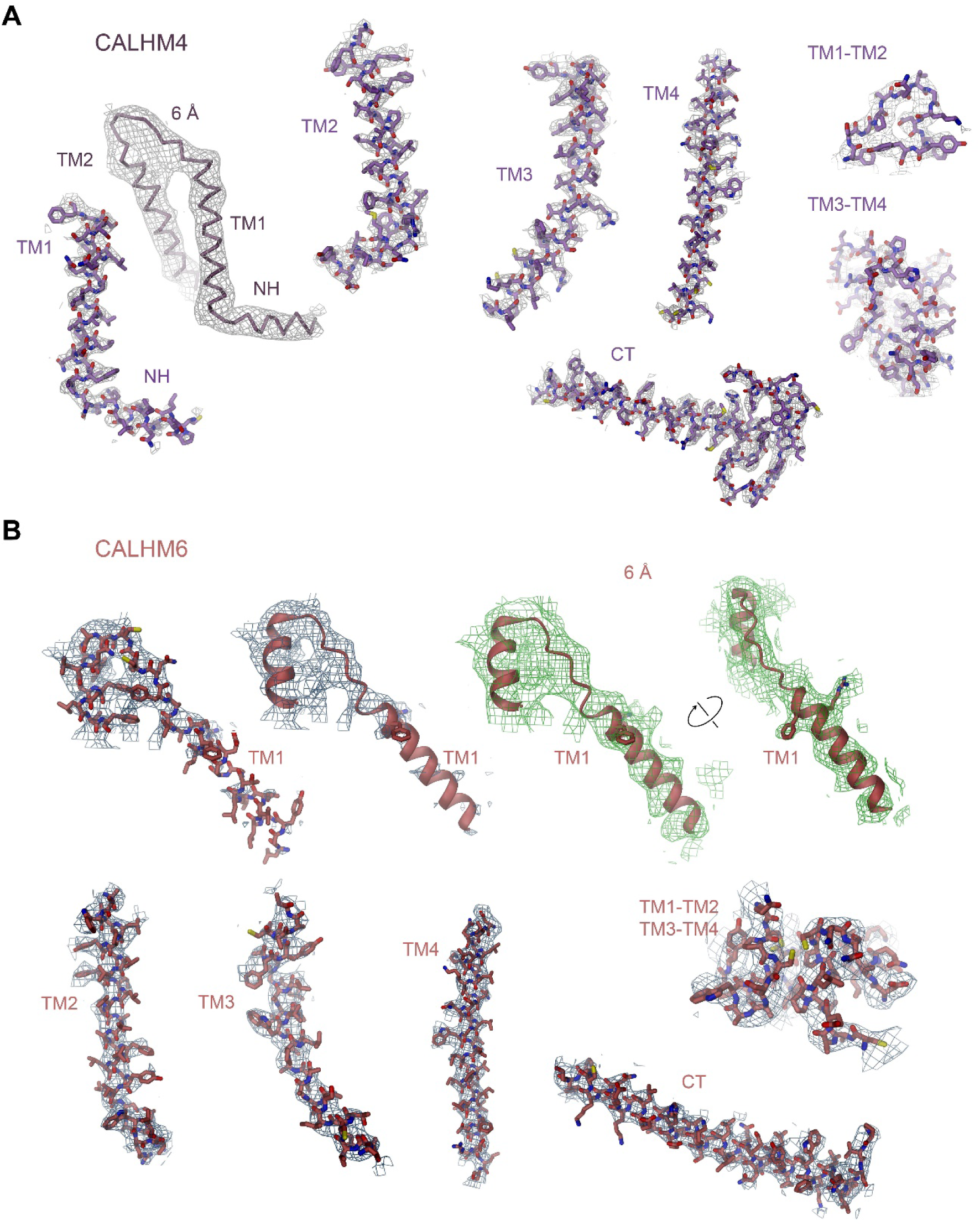
Cryo-EM density of CALHM4 and CALHM6. (**A**) Cryo-EM density at 3.9 Å of selected regions of the undecameric CALHM4 structure recorded from a sample obtained in the presence of Ca^2+^ superimposed on the model. Structural elements are indicated, ‘6 Å’ marks cryo-EM density low-pass filtered to 6 Å superimposed on a Cα trace of NH and TM1, which illustrates the helicity of the entire NH region that is partly not defined in the density at higher resolution due to its intrinsic mobility. (**B**) Cryo-EM density at 4.5 Å of selected regions of the decameric CALHM6 structure superimposed on the model. Structural elements are indicated. ‘6 Å’ marks cryo-EM density low-pass filtered to 6 Å superimposed on a ribbon of TM1 and parts of TM2, which displays extended density for the entire TM1 region that is partly not defined in the density at higher resolution due to its intrinsic mobility. Density is shown from two different views with indicated relationship. Large side chains that are recognizable in the low-pass filtered map are displayed as sticks.

**Figure 3—figure supplement 6.**
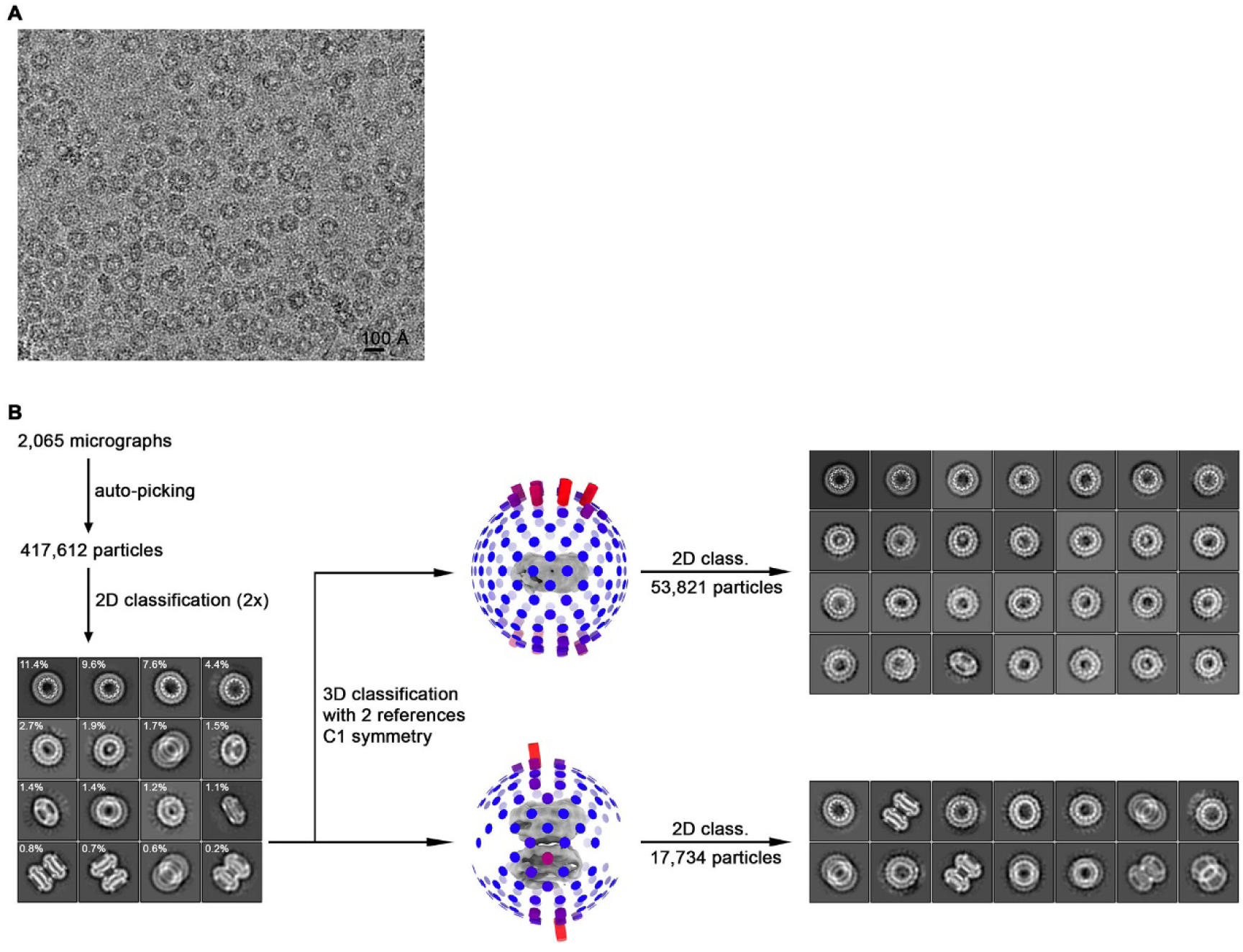
Cryo-EM reconstruction of CALHM2 in presence of Ca^2+^. (**A**) Representative cryo-EM micrograph acquired with a Tecnai G^2^ Polara microscope. (**B**) Data processing workflow. Non-symmetrized 3D classification with two reference models representing monomeric and dihedrally-related dimeric architectures as observed for CALHM6 and CALHM4, respectively, allowed to separate CALHM2 particles into two respective subsets. However, preferential orientation of the particles on a grid together with the presence of compositional heterogeneity in form of undecameric and dodecameric assemblies within each subset hindered generation of a high-resolution 3D reconstruction. 2D class averages calculated separately from monomeric and dimeric subsets highlight the predominance of views from the extracellular side.

**Figure 4—figure supplement 1.**
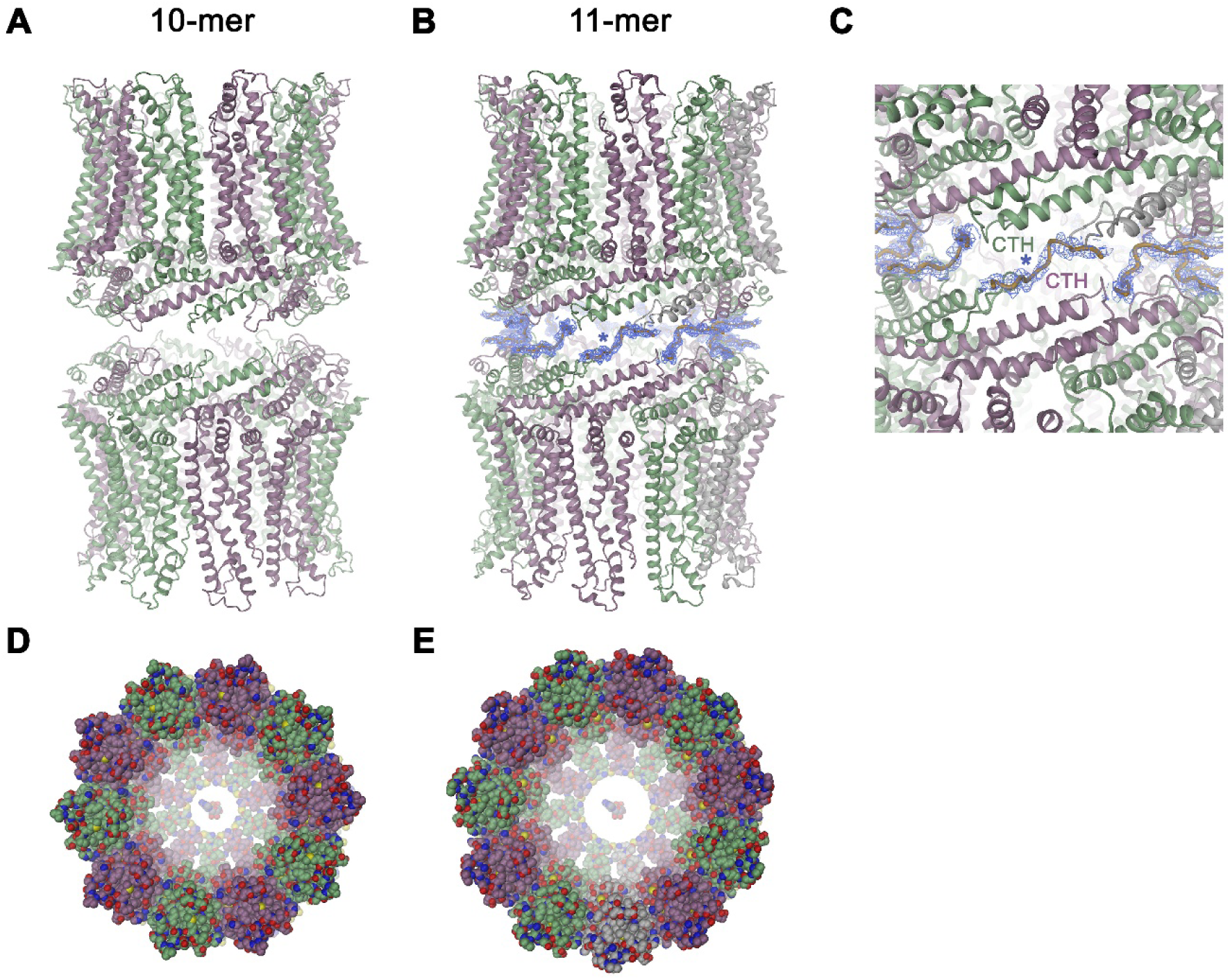
Features of the CALHM4 structure. Ribbon representation of (**A**), decameric and (**B**), undecameric channel pairs as observed in the CALHM4 dataset. In both cases, the proteins interact via their intracellular regions. Residual density at the C-terminus connecting both channels is shown as blue mesh and modeled as a poly-glycine chain. (**C**) Close-up of the interface between channel pairs shown in B. Space-filling models of the decameric (**D**), and undecameric (**E**) CALHM4 channel viewed from the extracellular side with ATP molecules shown for reference. Subunits are colored in lilac and green.

**Figure 5—figure supplement 1.**
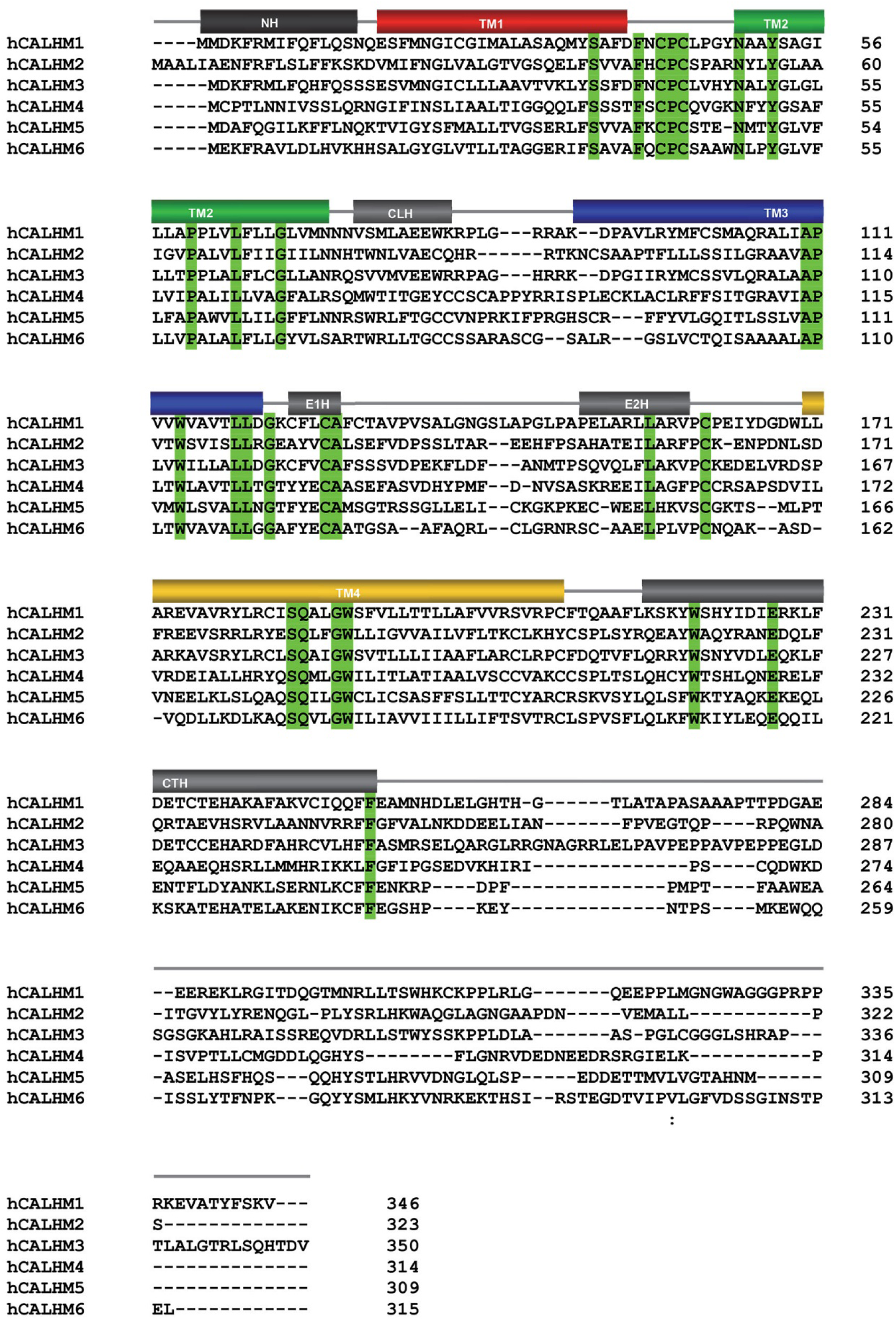
Sequence and topology. Sequence alignment of the human CALHM paralogs CALHM1 (NP_001001412.3), CALHM2 (NP_057000.2), CALHM3 (AAH43367.1), CALHM4 (NP_001353007.1), CALHM5 (NP_714922.1) and CALHM6 (NP_001010919.1). Identical residues are highlighted in green, secondary structure elements of CALHM4 are indicated above.

**Figure 5—figure supplement 2.**
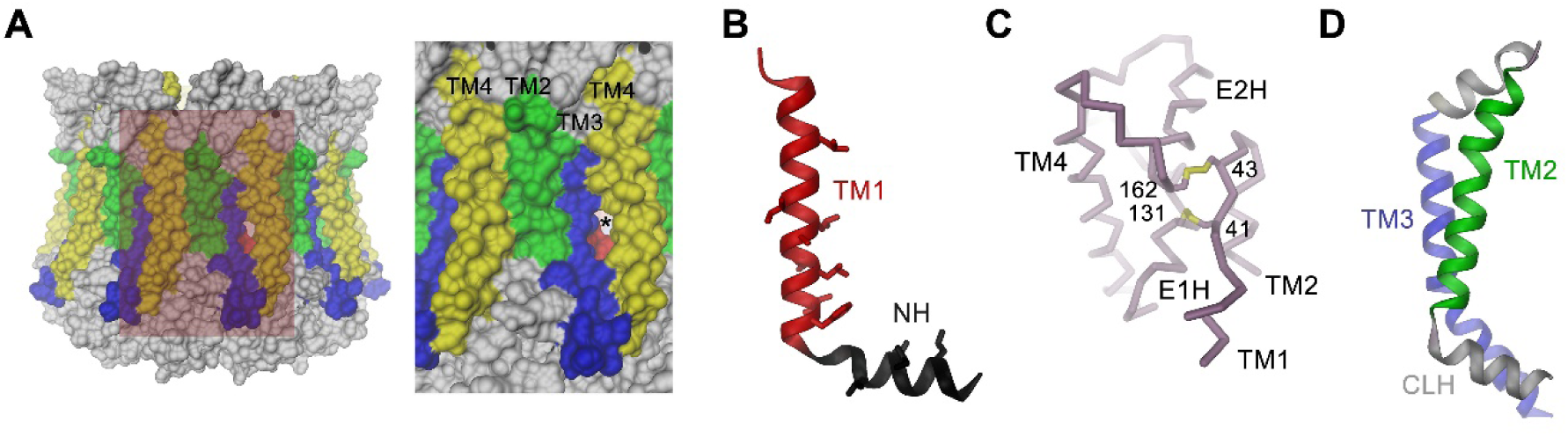
Features of the CALHM4 structure. (**A**) Molecular surface of the CALHM4 decamer viewed from within the membrane (left) and zoom into the highlighted region showing the interface between two subunits (right). Asterisk indicates location of a small fenestration on the surface between TM3 and TM4 of the same subunit. (**B**) Ribbon representation of TM1 and the N-terminal α-helix NH with side chains of hydrophobic residues facing the pore displayed as sticks. (**C**) Cα-trace of the extracellular region of CALHM4 with disulfide bridges shown as sticks and labeled. (**D**) Ribbon representation of TM2, the intracellular α-helix CLH and TM3. A, B, D, Transmembrane segments are colored as in Figure 5A and labeled.

**Figure 6—figure supplement 1.**
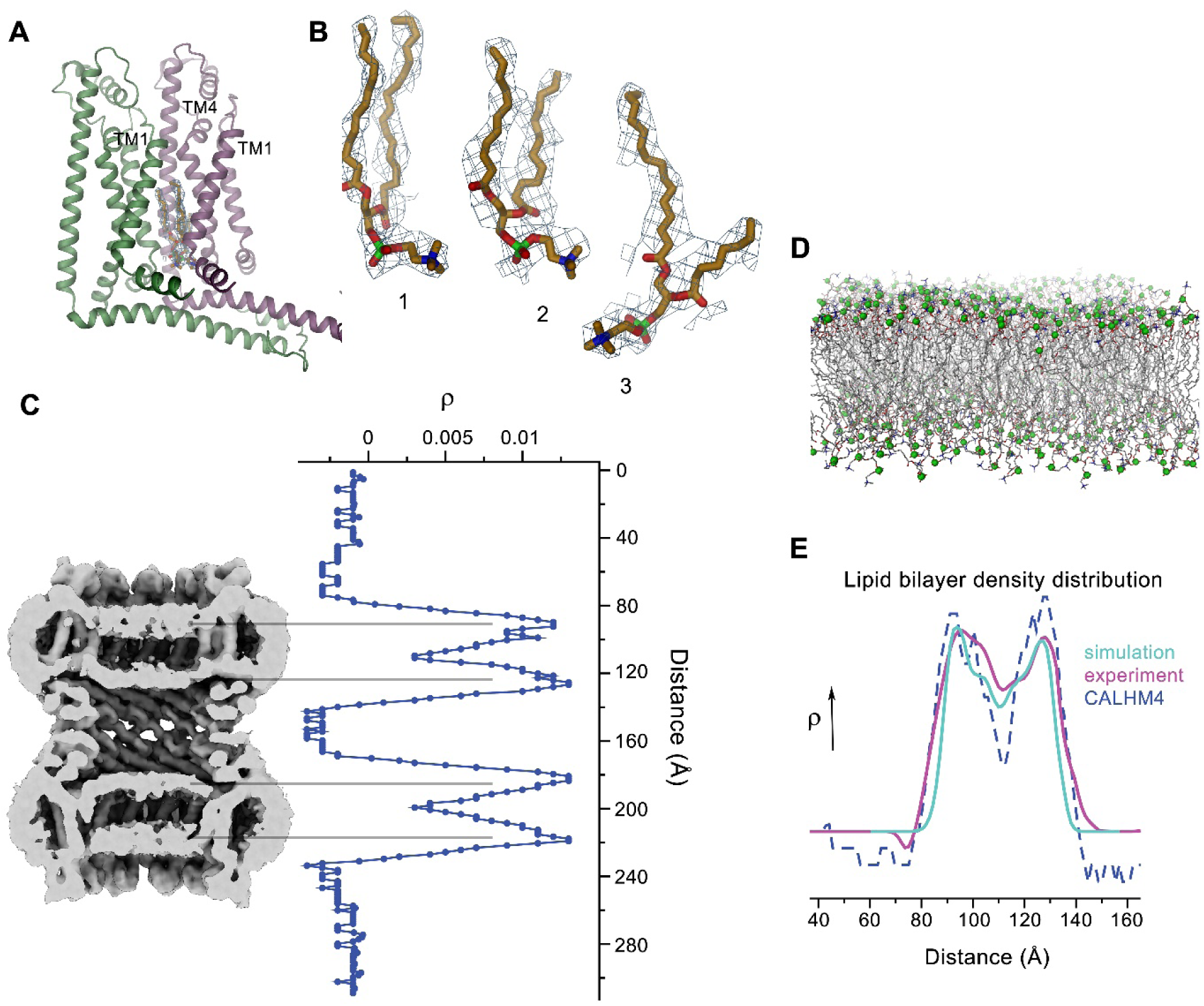
Lipid interactions in the pore of CALHM4. (**A**) Interface between α-helices TM1 of two neighboring subunits with two bound lipid molecules. View is similar as in Figure 6A. (**B**) Lipids modeled into residual electron density at the interface displayed in A. Lipids 2 and 3 could occupy the cavity at the same time. 1 shows alternative interpretation of the density with a single lipid. (**C**) Slice through the pore region of a pair of interacting CALHM4 decamers as observed in the CALHM4 data viewed from within the membrane. Shown is non-averaged density at low contour. The distribution (right) shows two regions of increased density corresponding to bilayers of either phospholipids or detergents in each channel with similar properties. The observed density distribution quantitatively matches equivalent distributions obtained from membrane simulations (**D**), shown as stick models with the phosphate atoms highlighted as green spheres and the cryo-EM density of liposomes (**E**).

**Figure 7—figure supplement 1.**
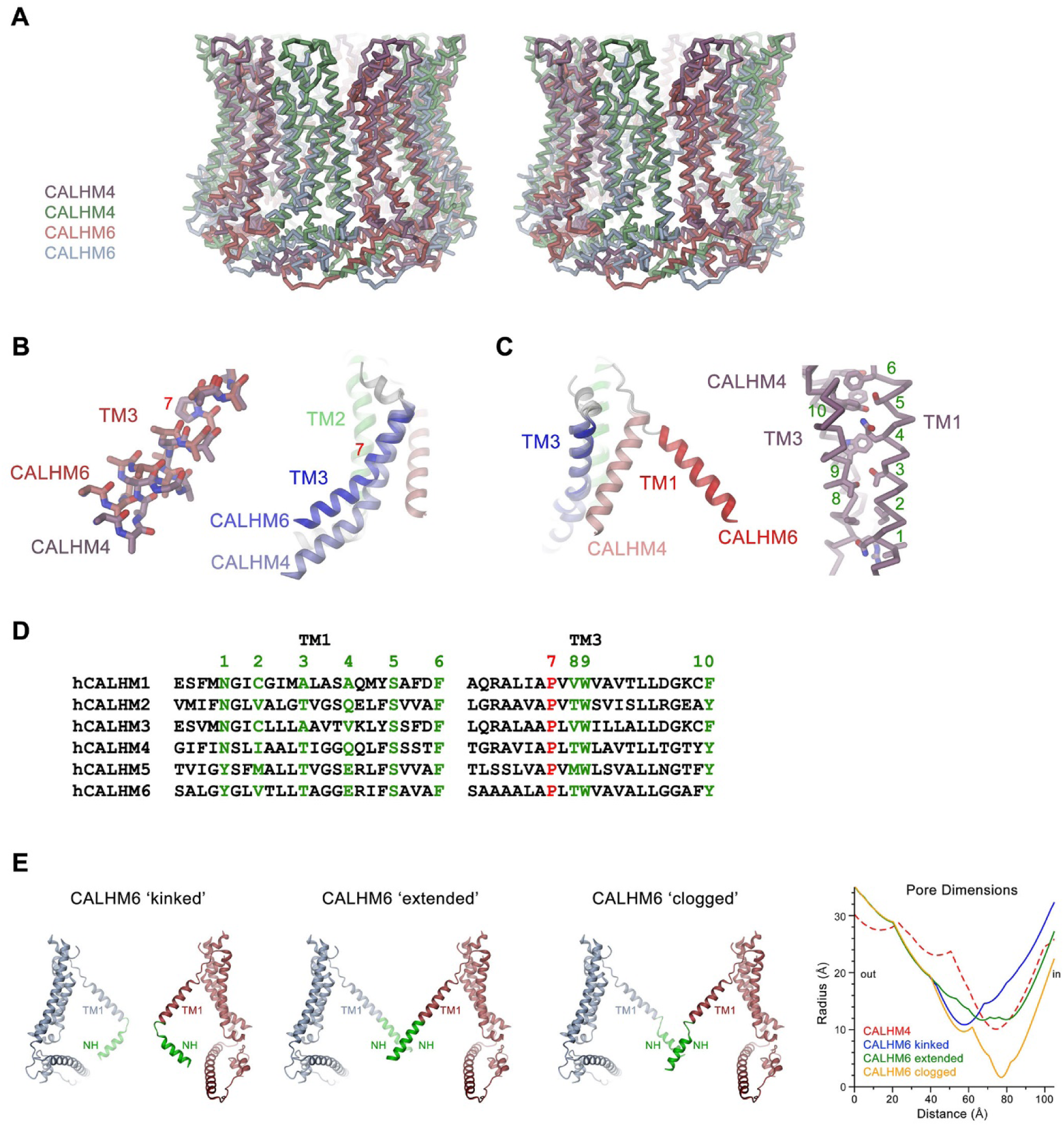
Features of the CALHM6 structure. (**A**) Stereo view of a superposition of the CALHM4 and CALHM6 decamers. The proteins are shown as Cα-traces. Coloring is as in Figures 4 and 6. (**B**) Close-up of the conformational change in TM3 from the cylindrical conformation displayed in CALHM4 to the conical conformation displayed in CALHM6. Right panel shows ribbon, left panel a stick model of TM3 with side chains truncated to their Cβ position except for a conserved proline at the site of conformational changes where the entire side chain is displayed. (**C**) Close-up of the conformational change of TM1 with molecules displayed as ribbons (left) and interaction region of TM1 and TM3 in the CALHM4 structure with the protein shown as Cα trace and side chains of interacting residues as sticks (right). B and C, coloring of ribbons is as in Figure 7D with secondary structure elements in CALHM4 shown in brighter shades of the same color. (**D**) Sequence alignments of TM1 and TM3 of different CALHM paralogs. The conserved proline displayed in B is colored in red, selected residues at the TM1 TM3 interaction interface displayed in C are colored in green. B, C, Numbering corresponds to positions labeled in D. (**E**) Hypothetical pore conformations of the conical conformation displayed in the CALHM6 structure. The disordered N-terminal helix NH (green) was modeled in different conformations. Left, ‘kinked’ conformation where the mutual relationship between TM1 and NH is as in CALHM4, center left, ‘extended’ conformation where NH has changed its conformation to extend the TM1 helix. Center right, ‘clogged’ conformation where the mobile NH helix has moved towards the pore axis. The 10-fold symmetry is maintained in the ‘kinked’ and ‘extended’ conformations and broken in the ‘clogged’ conformation. Opposite subunits of the decameric structures are displayed. Right, pore radii of the different modeled pore conformations of CALHM6 calculated with HOLE (Smart et al., 1996) compared to the CALHM4 decamer (red dashed) display similar constricting pore radius as for CALHM4 for ‘kinked’ and ‘extended’ conformations and an occluded pore in case of the ‘clogged’ conformation.

**Figure 8—figure supplement 1.**
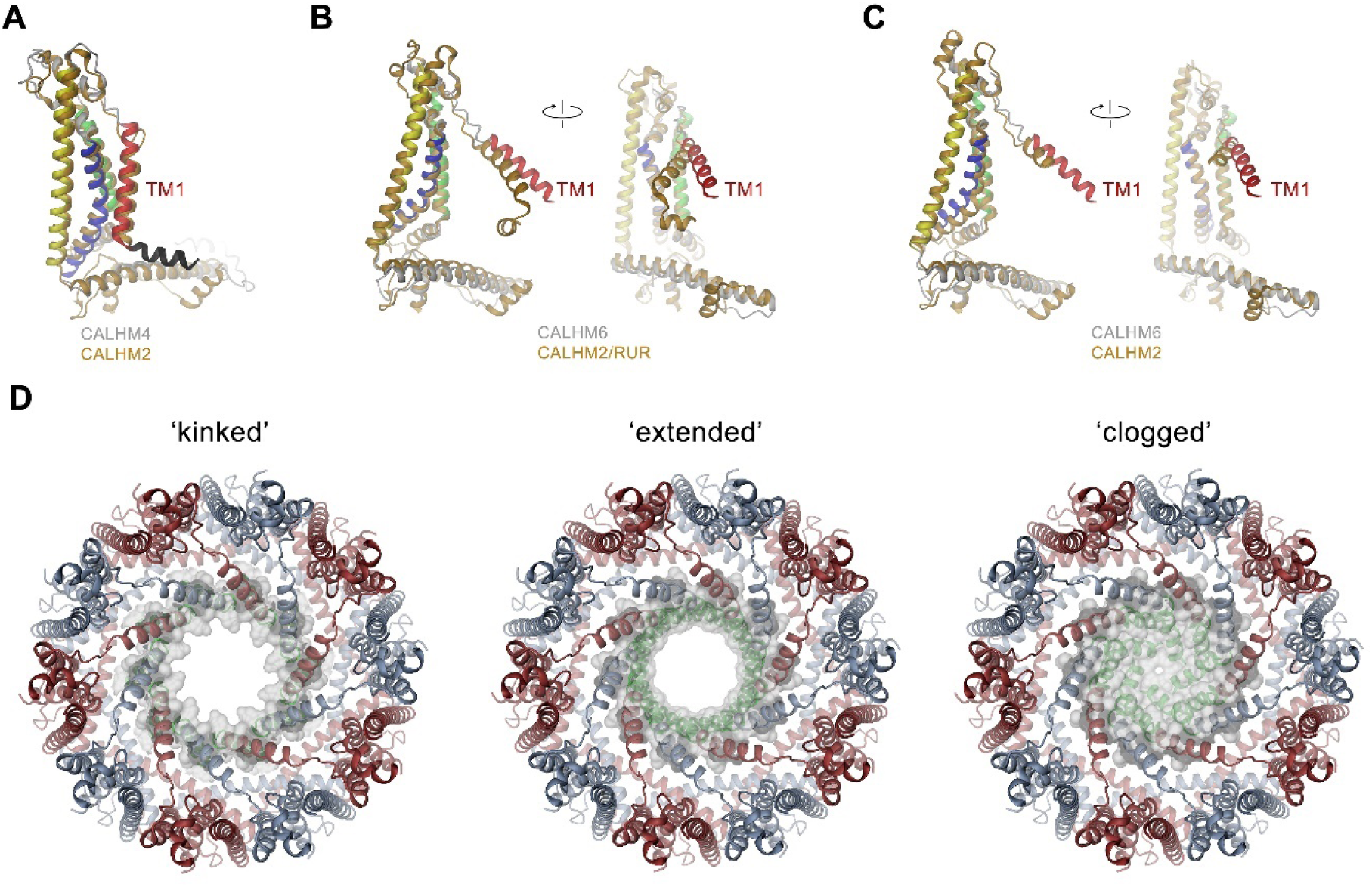
Comparison of pore conformations. (**A**) Comparison of the structures of subunits determined in this study to recent structures of CALHM2. Left, structure of the cylindrical conformation of CALHM4 superimposed on the equivalent conformation of CALHM2 (PDBID 6UIV). Superposition of the conical CALHM6 conformation with (**B**), the CALHM2 structure in complex with the putative inhibitor ruthenium red (RUR) (PDBID 6UIW) and (**C**), an equivalent conformation in the absence of RUR (PDBID 6UIX), both showing a different orientation of TM1. A-C, CALHM4 and 6 structures are colored as in Figure 5A, CALHM2 structures are colored in beige. B and C show two views of the subunit with indicated mutual relationship. (D) View of the three potential conformations of CALHM6 with modeled N-termini (green) as defined in Figure 7—figure supplement 1E. The molecular surface at the pore constriction is shown superimposed on the structure. A-D, proteins are displayed as ribbons.

**Video 1.** Cryo-EM density map of a single subunit of CALHM4. Shown is the cryo-EM map of the protein in detergent with the atomic model superimposed.

**Video 2.** Cryo-EM density map of a single subunit of CALHM6. Shown is the cryo-EM map of the protein in detergent with the atomic model superimposed.

**Video 3.** Morph between the cylindrical pore conformation of CALHM4 and the conical pore conformation displayed in the ‘extended’ model of CALHM6. In the latter the N-terminal α-helix NH is modeled in continuation of TM1 resulting in a constricting pore diameter similar to CALHM4.

